# ChIP-DIP: A multiplexed method for mapping hundreds of proteins to DNA uncovers diverse regulatory elements controlling gene expression

**DOI:** 10.1101/2023.12.14.571730

**Authors:** Andrew A. Perez, Isabel N. Goronzy, Mario R. Blanco, Jimmy K. Guo, Mitchell Guttman

**Affiliations:** Division of Biology and Bioengineering, California Institute of Technology, Pasadena CA 91125, USA; David Geffen School of Medicine, University of California, Los Angeles, Los Angeles, CA 90095, USA; Division of Chemistry and Chemical Engineering, California Institute of Technology, Pasadena, CA 91125, USA; Keck School of Medicine, University of Southern California, Los Angeles, CA 90089, USA

## Abstract

Gene expression is controlled by the dynamic localization of thousands of distinct regulatory proteins to precise regions of DNA. Understanding this cell-type specific process has been a goal of molecular biology for decades yet remains challenging because most current DNA-protein mapping methods study one protein at a time. To overcome this, we developed ChIP-DIP (ChIP Done In Parallel), a split-pool based method that enables simultaneous, genome-wide mapping of hundreds of diverse regulatory proteins in a single experiment. We demonstrate that ChIP-DIP generates highly accurate maps for all classes of DNA-associated proteins, including histone modifications, chromatin regulators, transcription factors, and RNA Polymerases. Using these data, we explore quantitative combinations of protein localization on genomic DNA to define distinct classes of regulatory elements and their functional activity. Our data demonstrate that ChIP-DIP enables the generation of ‘consortium level’, context-specific protein localization maps within any molecular biology lab.

## INTRODUCTION

Although every cell in the body contains the same genomic DNA sequence, distinct cell-types express different genes to enable cell-type specific function. Cell-type specific gene regulation involves the coordinated activity of thousands of regulatory proteins that localize at precise DNA regions to activate, repress, and quantitatively control levels of transcription. Each genomic DNA region is organized around nucleosomes^1^, which contain histone proteins that undergo extensive post-translational modifications^2,3^ and together define cell-type specific chromatin states. Chromatin state is controlled by chromatin regulators that directly read, write and erase specific histone modifications^4,5^, as well as control nucleosome positioning and DNA accessibility^6,7^. This ultimately determines which genomic regions are accessible for binding by sequence-specific transcription factors^8^, the enzymes that transcribe DNA into RNA (RNA polymerases)^9^ and other general and specific regulatory proteins that promote or suppress transcriptional initiation^10,11^.

Understanding how regulatory protein binding gives rise to cell type-specific gene expression has been a central goal of molecular biology for decades^4^. Over the past 20 years, significant technical advances have enabled genome-wide mapping of regulatory proteins and histone modifications (e.g. ChIP-Seq)^12–15^, improved binding site resolution (ChIP-Exo)^16,17^, increased sample throughput (e.g. through automation and/or sample pooling)^18,19^, and mapping within a limited numbers of cells (e.g. CUT&RUN/CUT&Tag)^20–22^. Yet, while these innovations have enabled new applications and uncovered critical new insights into gene regulation, all of these approaches still work by studying a single protein at a time. The two exceptions are multiplexed Cut&Tag^23,24^ and MAbID^25^, which can measure up to three or six histone modifications, but not transcription factors or other regulatory proteins, in a single experiment^24^. Due to the large number of distinct regulatory proteins and histone modifications involved and the cell-type specific nature of their regulatory interactions, this one-at-a-time mapping approach makes it extremely difficult to construct a comprehensive understanding of gene regulation.

Initial attempts to overcome this challenge led to the formation of various international consortia that aimed to generate reference maps of hundreds of proteins within a small number of cell types (ENCODE^26^, PsychENCODE^27^, ImmGen^28^, etc.). Although these efforts have provided many critical insights^29–31^, because protein binding maps and gene expression programs are intrinsically cell type-specific^32,33^, it is not possible to study cell type-specific regulation using maps generated from reference cell lines^34^. To date, most mammalian cell types, experimental and disease models, and model organisms remain uncharacterized. Generating additional cell type-specific regulatory maps currently requires consortium-level effort (dozens of labs across the world), time (many years), and resources (>$100 million) for each biological system. Accordingly, there is a clear need for a highly scalable, multiplexed protein profiling method that can increase throughput of protein mapping by orders of magnitude and profile the diverse categories of DNA-associated proteins, including classes that have been traditionally easier to map (e.g. histone modifications) and those that have been more challenging (e.g. transcription factors)^35^. Such a method would allow any lab to generate comprehensive maps for any cell type of interest in a rapid and cost-effective way and would enable exploration of key questions that is not currently possible.

To address this need, we developed chromatin immunoprecipitation done-in-parallel (ChIP-DIP), a scalable platform that enables simultaneous, genome-wide mapping of hundreds of diverse regulatory proteins within a single experiment. We utilized ChIP-DIP to generate data from ∼180 distinct DNA associated proteins in human or mouse cells, including a single multiplexed ChIP-DIP experiment containing >225 different antibodies targeting ∼160 distinct proteins (**Supplemental Table 1**). We show that ChIP-DIP generates accurate genome-wide maps, equivalent to those generated by traditional approaches, that are highly robust regardless of the number of antibodies or protein composition contained within a pool and across a range of input cell numbers. We show that ChIP-DIP enables accurate mapping of all classes of DNA-associated proteins, including histone modifications, chromatin regulators, transcription factors and other sequence-specific DNA binding proteins, and RNA Polymerases. Together, our results demonstrate that ChIP-DIP enables the generation of ‘consortium level’ comprehensive, context-specific protein localization maps within any experimental system and within any molecular biology lab and enables the exploration of complex, combinatorial patterns of protein localization that define regulatory activity. Beyond the immediate applications, ChIP-DIP can be directly integrated into a suite of existing split-pool approaches to enable highly multiplexed mapping of protein localization within single cells in combination with measurements of 3D structure^36^, ncRNA localization, and nascent transcription^37^.

## RESULTS

### ChIP-DIP: A highly multiplexed method for mapping DNA-associated proteins

To enable highly multiplexed, genome-wide mapping of hundreds of DNA-associated proteins in a single experiment, we developed ChIP-DIP (ChIP Done In Parallel) (**Figure 1A)**. ChIP-DIP works by (i) using a rapid, modular, and simple approach to couple individual antibodies to beads containing a unique oligonucleotide tag (**Figure S1A**), (ii) combining sets of different antibody-bead-oligo conjugates to create an antibody-bead pool, (iii) performing ChIP using this pool, (iv) conducting split-and-pool barcoding to match antibody-bead-oligo conjugates to specific genomic DNA regions^38–40^, and (v) sequencing DNA and computationally matching split-pool barcodes that are shared between genomic DNA and the antibody tag. We refer to all unique reads containing the same split-pool barcode as a cluster. We combine DNA reads from all clusters corresponding to the same antibody to generate a protein localization map for each individual protein. The output of a ChIP-DIP experiment is analogous to the data generated in a traditional ChIP-Seq experiment, however instead of a single map, ChIP-DIP provides a set of distinct maps – one for each antibody utilized (**Figure 1B**).

**Figure 1:**
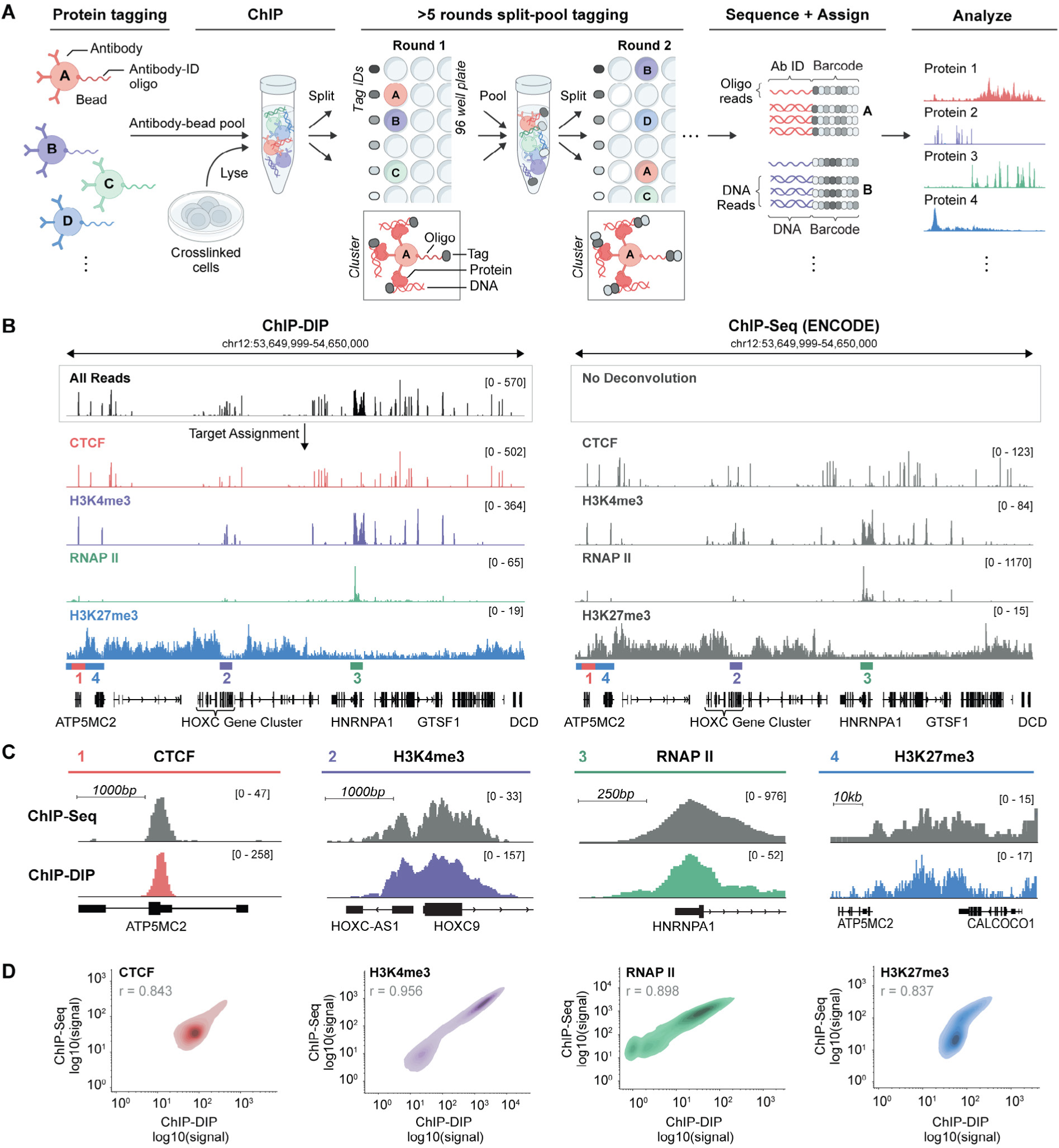
ChIP-DIP: A highly multiplexed method for mapping proteins to genomic DNA. **(A)** Schematic of the ChIP-DIP method. Beads are coupled with an antibody and associated oligonucleotide (antibody-ID). Sets of beads are then mixed (antibody-bead pool) and used to perform ChIP. Multiple rounds of split-and-pool barcoding are performed to identify molecules bound by the same Protein G bead. DNA is sequenced and genomic DNA and antibody oligos containing the same split-and-pool barcode are grouped into a cluster, which are used to assign genomic DNA regions to their linked antibodies. All DNA reads corresponding to the same antibody are used to generate protein-localization maps. **(B)** Protein localization maps over a specific human genomic region (hg38, chr12:53,649,999-54,650,000) for four protein targets - CTCF, H3K4me3, RNAP II and H3K27me3. Left panel: Protein localization generated by ChIP-DIP in K562. Top track shows read coverage prior to protein assignment and bottom four tracks correspond to read coverage after assignment to individual proteins. Right panel: ChIP-Seq data generated by ENCODE within K562 for these same 4 proteins are shown for the same region. To enable direct comparison of scales between datasets, we normalized the scale to coverage per million aligned reads. Scale is shown from 0 to maximum coverage within each region. **(C)** Comparison of ChIP-DIP and ChIP-Seq maps over specific regions corresponding to zoom-ins of the larger region shown in (B). The locations presented are demarcated by colored bars above the gene track of (B). Scale shown similar to (B). **(D)** Genome-wide comparison (density plots of signal correlation) between the localization of each individual protein measured by ChIP-DIP (x-axis) or ChIP-Seq (y-axis). Points are measured genome-wide across 10kb windows (CTCF, H3K27me3) or all promoter intervals (H3K4me3, RNAP II).

To ensure that chromatin-antibody-bead-oligo conjugates remain intact throughout the ChIP-DIP procedure (rather than dissociating and reforming new complexes), we designed a series of experiments to measure dissociation between (i) oligo and bead, (ii) antibody and bead, or (iii) antibody and chromatin (**Figure S1B, Supplemental Note 1**). We observed minimal dissociation for any of these cases; most beads contain a single oligo type (>95%, **Figure S1C**), beads without a coupled antibody are associated with minimal chromatin (<0.5%, **Figure S1D**) and most purified chromatin originates from the initial capture (> 94%, **Figure S1E**).

To test whether ChIP-DIP can accurately map genome-wide protein localization, we performed a ChIP-DIP experiment in human K562 cells using four well-studied proteins: (1) the CTCF sequence-specific DNA binding protein that binds to insulator sequences^41^, (2) the H3K4me3 histone modification that localizes at the promoters of active genes^42,43^, (3) the RNA Polymerase II enzyme that transcribes RNA^44^ and (4) the H3K27me3 histone modification that accumulates over broad genomic regions that are associated with polycomb-mediated transcriptional repression^42,43^.

We observed localization patterns that are highly comparable at specific genomic sites (**Figure 1B-C**) and highly correlated genome-wide (r=0.837-0.956, **Figure 1D**) to ChIP-Seq profiles generated by the ENCODE consortium^45–47^ (**Supplemental Table 2**). These genome-wide profiles were highly consistent even when using antibody pools containing different numbers of antibodies and protein composition (**Figure 2A-D, Supplemental Table 3**), including pools containing independent antibodies targeting the same protein (CTCF) or multiple proteins within the same complex (e.g., members of the PRC1/2 complex, **Figure S2**). Because ChIP-DIP enables simultaneous mapping of many proteins within the same experiment, we reasoned that it may dramatically reduce the total number of input cells required per experiment^48^ (**Methods**). Indeed, we observed strong genome-wide correlations and peak overlap when measuring these same proteins across a range of input cell numbers (**Figure 2E-H, Figure S3, Supplemental Table 4**). Because ChIP-DIP can generate dozens of individual maps from the same lysate, this further reduces the effective number of cells required for each protein target. In this example, we observed strong correlations with maps generated from ∼1,400 cells per protein. Together, these results demonstrate that ChIP-DIP is highly robust and generates data that are highly comparable to those generated by standard methods.

**Figure 2:**
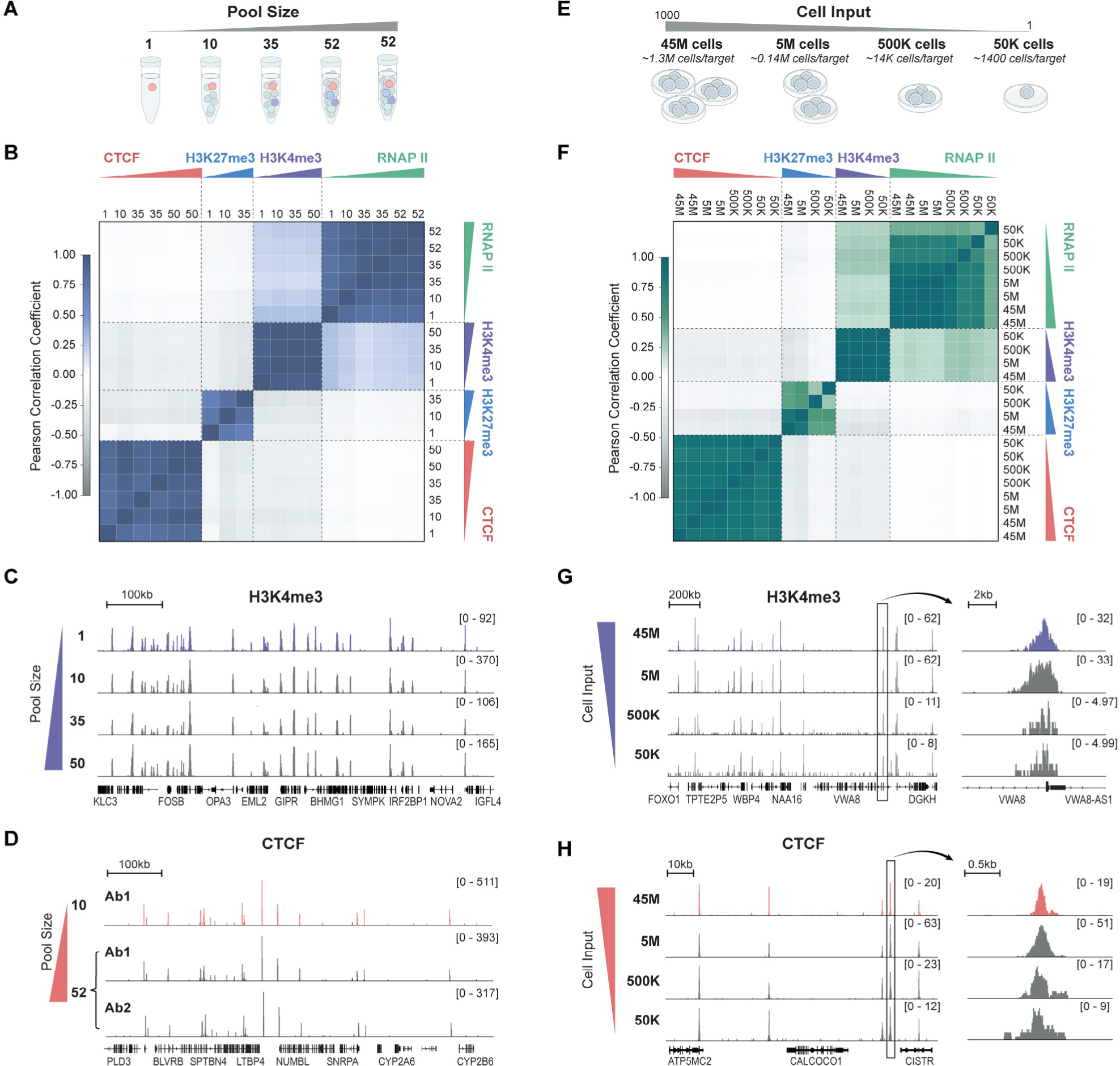
ChIP-DIP accurately maps large sets of proteins using low-levels of cell lysate. **(A)** Schematic of experimental design to test scalability of antibody-bead pool size and composition. **(B)** Correlation heatmap for protein localization maps of four proteins – CTCF, H3K4me3, RNAP II and H3K27me3 – generated using antibody pools of four different sizes and compositions (see Methods). Pool sizes are listed along top and left axis. Replicate proteins in the same pool indicate that a different antibody was used for that protein. Some antibodies were not included in every pool. **(C)** Comparison of H3K4me3 localization over a specific genomic region (hg38, chr19:45,345,500-46,045,500) when measured within various antibody pool sizes and compositions. **(D)** Comparison of CTCF localization over a specific genomic region (hg38, chr19:40,349,999-41,050,000) when measured within a pool of 10 antibodies containing a single CTCF-targeting antibody (top) or two different CTCF-targeting antibodies within a pool of 52 antibodies (bottom). **(E)** Schematic of experimental design to test the amount of cell input required for ChIP-DIP. **(F)** Correlation heatmap for protein localization maps of four targets – CTCF, H3K4me3, RNAP II and H3K27me3 – generated using various amounts of input cell lysate (see Methods). Amounts of input cell lysate are listed along top and left axis. **(G)** Comparison of H3K4me3 localization over a specific genomic region (hg38, chr13:40,600,000-42,300,000) when measured using various amounts of input cell lysate. **(H)** Comparison of CTCF localization over a specific genomic region (hg38, chr12:53,664,000-53,764,000) when measured using various amounts of input cell lysate.

### ChIP-DIP accurately maps hundreds of diverse DNA-associated proteins

We next explored whether ChIP-DIP can simultaneously map proteins from distinct categories, some of which have been traditionally easier to map than others^49,50^. To do this, we performed ChIP-DIP on >60 distinct proteins in human K562 cells and >160 distinct proteins in mouse embryonic stem cells (mESCs) across six experiments (**Supplemental Table 1**). These included 39 histone modifications (HMs), 67 chromatin regulators (CRs), 51 transcription factors (TFs), and all three RNA Polymerases (RNAPs) and 4 of their modified forms.

#### Histone modifications

Histone modifications define cell-type specific chromatin states and have proven incredibly useful for annotating cell type-specific regulatory elements^51^. We mapped 39 histone modifications – including 18 acetylation, 17 methylation, 3 ubiquitination and 1 phosphorylation marks – in either mESCs or K562s (**Figure 3A**). We confirmed the localization of five histone modifications commonly used to demarcate five functional states^52^, as well as additional modifications associated with each state (**Figure S4A-F**): enhancer regions^53^ (H3K4me1, H3K4me2, H3K27ac, **Figure 3B**), transcribed regions^42,54,55^ (H3K36me3, H3K79me1/2, **Figure 3C**), promoter regions^42,43,56^ (H3K4me3, H3K9ac, **Figure 3D**), polycomb-repressed regions^57^ (H3K27me3, H2AK119ub, **Figure 3E**), and constitutive heterochromatin regions^58^ (H3K9me3, H4K20me3, **Figure 3F**). These data indicate that ChIP-DIP accurately maps histone modifications with distinct genome-wide patterns (broad and focal localization), that represent distinct activity states (active and repressive), and that localize at distinct functional elements (promoters, enhancers, gene bodies, and intergenic regions).

**Figure 3:**
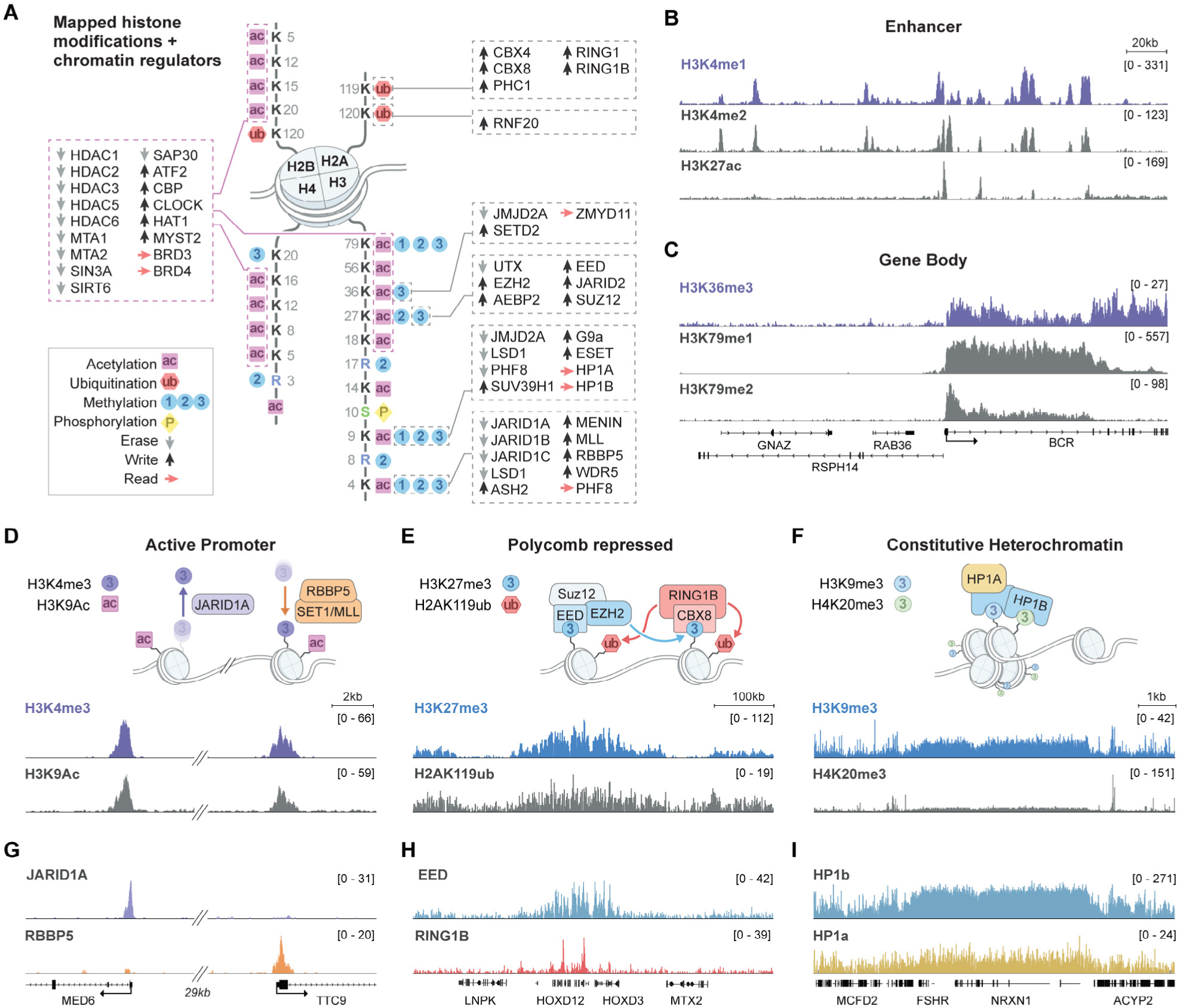
ChIP-DIP accurately maps dozens of functionally diverse histone modifications and chromatin regulators. **(A)** Illustration of the diverse histone modifications and chromatin regulatory proteins mapped in K562 or mESC using ChIP-DIP. **(B-C)** Visualization of multiple histone modifications across a genomic region (hg38, chr22:23,050,000-23,290,000) in K562 corresponding to multiple histone modifications associated with **(B)** enhancers – H3K4me1, H3K4me2 and H3K27Ac and **(C)** active gene bodies – H3K36me3, H3K79me1 and H3K79me2. **(D)** Top: Schematic of histone modifications and chromatin regulators associated with active promoters. Bottom: Visualization of multiple histone modifications associated with active promoters - H3K4me3 and H3K9Ac – across a genomic region (mm10, chr12:81,590,000-81,636,000) in mouse ESCs. Hashmarks indicate an intervening 29kb region that is not shown. **(E)** Top: Schematic of histone modifications and chromatin regulators associated with polycomb-mediated repression. Bottom: Visualization of multiple histone modifications associated with polycomb-mediated repression – H3K27me3 and H2A119ub – across a genomic region (hg38, chr2:175,846,000-176,446,000) containing the silenced HOXD cluster in K562. **(F)** Top: Schematic of histone modifications and chromatin regulators associated with constitutive heterochromatin. Bottom: Visualization of multiple histone modifications associated with constitutive heterochromatin – H3K9me3 and H4K20me3 – across a genomic region (hg38, chr2:46,200,000-55,700,000) in K562. **(G)** Visualization of an H3K4me3-associated eraser (JARID1A) and writer component (RBBP5) across the same genomic region as (D). **(H)** Visualization of PRC2 (EED) and PRC1 (RING1B) components across the same genomic region as (E). **(I)** Visualization of HP1b and HP1a across the same genomic region as (F).

#### Chromatin regulators

Chromatin regulators (CRs) are responsible for reading, writing, and erasing specific histone modifications and are critical for the establishment, maintenance, and transition between chromatin states^59,60^. We measured 67 CRs associated with various histone methylation, acetylation, and ubiquitination marks, as well as with DNA methylation, in either mouse ES or human K562 cells (**Figure 3A**). As expected, we observe that an eraser (JARID1A)^61^ and a writer (RBBP5-containing complex)^62^ of H3K4me3 localize at H3K4me3-modified promoter sites (**Figure3G**, **Figure S4G**). Additionally, we observed that components of the PRC1 (RING1B, CBX8)^63^ and PRC2 complex (EED, SUZ12, EZH2)^64^ co-localize and are enriched over genomic regions containing their respective histone modifications (H2AK119ub and H3K27me3, **Figure 3H**, **Figure S4H**). Similarly, we observed co-localization of two members of the Heterochromatin Protein 1 (HP1) family, HP1α and HP1ß, at genomic DNA regions containing their associated heterochromatin marks, H3K9me3 and H4K20me3^65^ (**Figure 3I**, **Figure S4I**). These data indicate that ChIP-DIP accurately maps chromatin regulators from diverse complexes and with distinct functional properties (i.e., modification recognition, enzymatic activity, chromatin packaging).

#### Transcription factors

Transcription factors (TFs) bind *cis*-regulatory elements in combinatorial patterns to control gene expression. Generating comprehensive maps of TF localization has proven difficult because there are large numbers of distinct TFs, most are cell type-specific, and they are challenging to map by ChIP-Seq because they tend to be lower in abundance and only transiently associated with DNA^66,67^. To explore whether ChIP-DIP can map large sets of TFs, we measured 15 TFs in K562 and 43 TFs in mESC, including constitutive (e.g. SP1 and USF2)^68,69^, stimulus-dependent (e.g. p53 and NRF1)^70–73^, and developmental/cell type-specific (e.g. Nanog and RFX1)^74,75^ DNA binding proteins^76^ (**Figure 4A**). We obtained high resolution binding maps for TFs in both cell types, with previously characterized TFs showing localization at their expected genomic DNA targets^68,71,73,77–79^ (**Figure 4A-B**, **Supplemental Table 5**). Using these genome-wide localization data, we accurately identify DNA binding motifs for each TF, including the 20bp dimer motif of p53^80^ and the 21bp RE-1 consensus sequence of REST^81^ (**Figure 4C**). Together, these data indicate that ChIP-DIP generates accurate, high-resolution binding maps of diverse TFs in multiple cell types.

**Figure 4:**
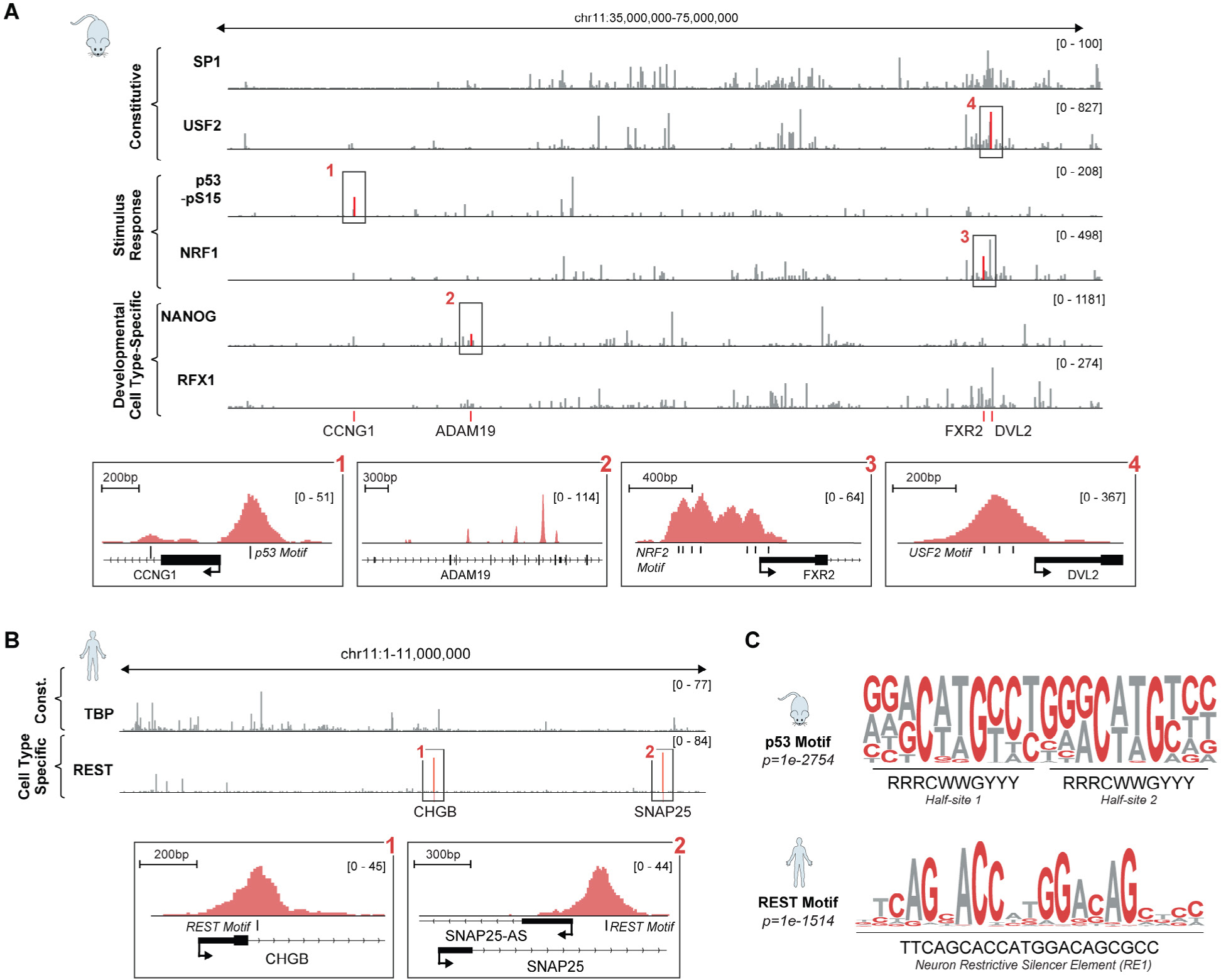
ChIP-DIP accurately maps dozens of transcription factors representing diverse functional classes. **(A)** Top: Visualization of six transcription factors (SP1, USF2, p53-pSer15, NRF1, NANOG, RFX1) representing three broad functional classes (constitutive, stimulus-response, development/cell type-specific) across a genomic region (mm10, chr11:35,000,000-75,000,000) in mESC. Bottom: Higher-resolution zoom-ins showing individual TF binding patterns at selected targets and motif localization as appropriate. (1) p53 binding the p53 response element on the Cyclin G1 gene promoter. (2) Nanog binding a cluster of sites internal to the developmental gene ADAM19. (3) Nuclear Respiratory Factor 1 (NRF1) binding multiple copies of its motif at the promoter of FXR2. (4) The constitutively active USF2 binding its triplicate E-box motif. **(B)** Visualization of TBP (constitutive) and REST (cell type-specific) across a genomic region (hg38, chr11:1-11,000,000) in K562 cells. Bottom: Higher-resolution zoom-ins highlight two individual peaks of RE-1 Silencing Transcription Factor/ Neuron-Restrictive Silencer Factor (REST/NRSF) at motif sites near promoters of known neuronal gene targets. **(C)** *de novo* generated motifs for p53 (top) in mESCs and REST (bottom) in K562 cells using binding sites identified using ChIP-DIP.

#### RNA Polymerases (RNAPs)

Different classes of RNA are transcribed by distinct RNA polymerases: RNA Polymerase I (RNAP I) transcribes the 45S ribosomal RNA (rRNA) encoding the 18S, 28S, and 5.8S rRNAs; RNAP II transcribes messenger RNAs and various non-coding RNAs, including snRNAs, snoRNAs and lncRNAs; and RNAP III transcribes diverse small RNAs, including transfer RNAs, 5S rRNA, 7SL, 7SK, and U6 snRNA^82^. We leveraged the power of ChIP-DIP to simultaneously map all three RNAPs and the post-translationally modified forms of RNAP II. We observed that each RNAP localizes with high selectivity to its corresponding classes of genes; RNAP I binds at rDNA, RNAP II at mRNA and snRNA genes, and RNAP III at tRNA genes (**Figure 5A**, **Figure S5A**). Moreover, we observed distinct localization patterns of different RNAP II phosphorylation states: serine 5 phosphorylated RNAP II localizes at promoters, while serine 2 phosphorylated RNAP II accumulates over the gene body and past the 3’ end of the gene (**Figure S5B-C**). These data indicate that ChIP-DIP accurately maps the localization of the three RNA polymerases – including multiple functional phosphorylation states of RNAP II – at distinct gene classes and gene features.

**Figure 5:**
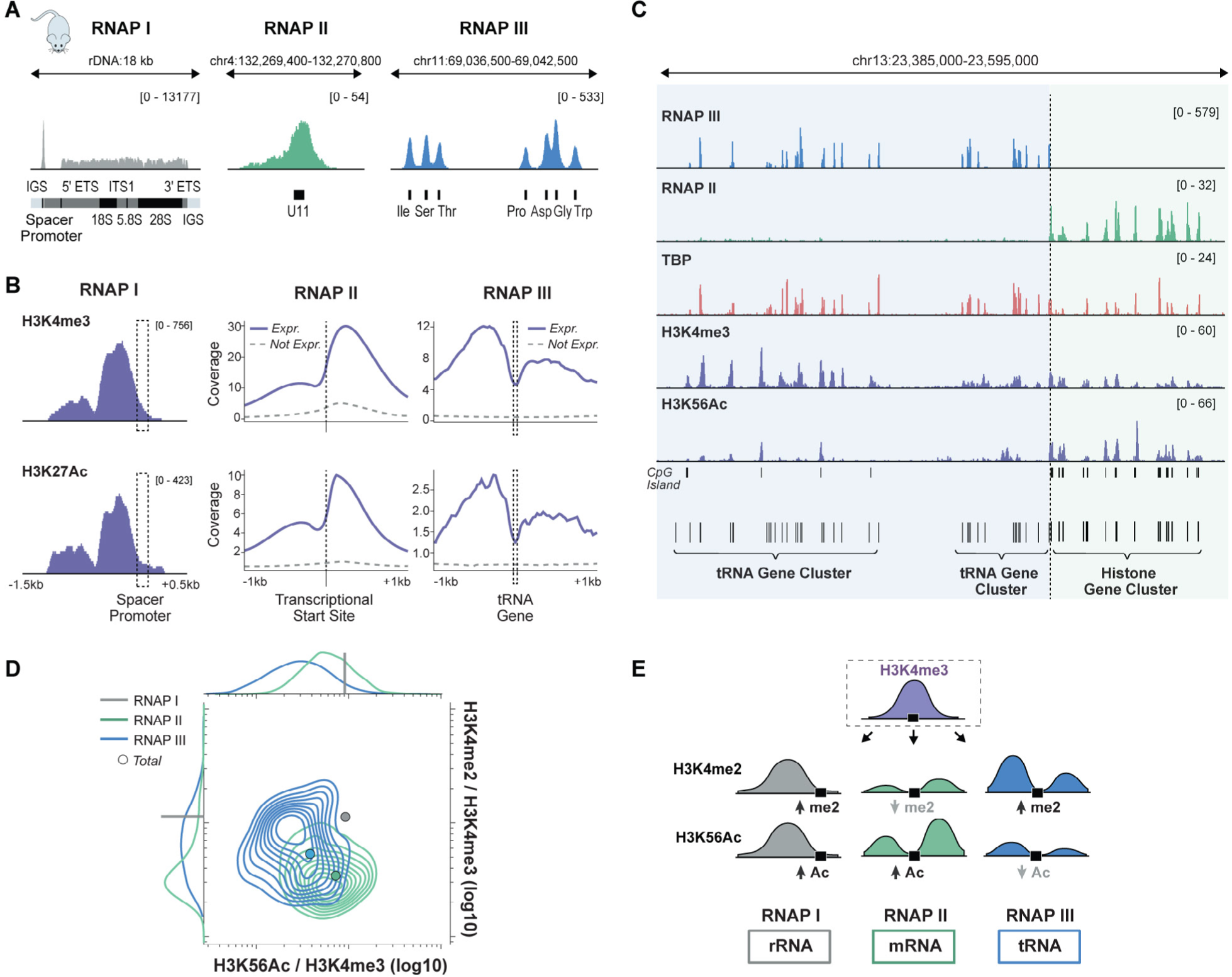
Distinct chromatin signatures define the promoters of each RNA Polymerase. **(A)** Visualization of RNAP I at the promoter and along the gene body of rDNA (left), RNAP II at a snRNA gene (middle), and RNAP III at a cluster of tRNA genes (right) in mouse ESCs. **(B)** Comparison of H3K4me3 and H3K27Ac profiles at the promoters of RNAP I, II and III genes. RNAP I is displayed over the rDNA spacer promoter (left) while RNAP II and III are displayed as metaplots across active (blue) and inactive (dashed gray) promoters. **(C)** Visualization of RNAP II and RNAP III along with the shared transcription factor TBP, and histone modifications H3K4me3 and H3K56Ac across a genomic region (mm10, chr13:23,385,000-23,595,000) containing a tRNA gene cluster (RNAP III transcribed genes) adjacent to a histone gene cluster (RNAP II transcribed genes), separated by dashed line. **(D)** Density distribution of H3K4me2/H3K4me3 versus H3K56Ac/H3K4me3 ratios at RNAP I, active RNAP II and active RNAP III promoters. Points show ratios when computed using the total sum of histone coverage over all promoters. Marginal distributions are shown for RNAP II and III along x and y-axis. Axes are log10 scaled. **(E)** Schematic showing relative levels of histone modifications H3K4me2 and H3K56Ac at H3K4me3 enriched regions and the associated RNAP promoter.

Together, these results establish ChIP-DIP as a modular, highly-multiplexed method that generates high-quality maps for a wide range of DNA-associated proteins spanning diverse biological functions.

### Integrated analysis of protein localization identifies functionally distinct *cis*-regulatory elements

Previous large-scale analyses have identified histone modifications that demarcate distinct genomic elements (e.g. promoters, enhancers, transcribed regions, etc.)^83^, their activity state (active, inactive, repressed), and regulatory potential (poised/primed for activation)^84^. However, because of the large number of histone modifications and regulatory proteins, many efforts have focused on mapping only five histone modifications to identify genomic features and regulatory states (i.e., H3K4me3, H3K4me1, H3K36me3, H3K9me3, and H3K27me3 marking promoters, enhancers, elongated transcripts, heterochromatin, and polycomb-mediated silencing, respectively)^52^. Because ChIP-DIP can map large numbers of diverse proteins, we asked whether additional combinations of histone modifications and regulatory proteins can provide additional information about distinct types, activity states, and regulatory potentials of *cis*-regulatory elements (promoters or enhancers) beyond those captured by the five commonly studied individual histone modifications.

### Promoter type and activity state are defined by combinations of histone modifications

H3K4me3 is generally thought to mark the promoters of actively transcribed RNAP II transcripts^42,43,85^. Consistent with this, we find H3K4me3 over the promoters of actively transcribed RNAP II genes, yet we also observe this modification near RNAP I promoters (ribosomal RNA) and many active RNAP III genes (tRNAs) (**Figure 5B**). Similarly, we observe that other histone modifications associated with active RNAP II promoters, including H3K4me2, H3K9Ac, H3K27Ac, and H3K56Ac, are also enriched at RNAP I and III genes (**Figure 5B-C, Figure S5D**). For example, focusing on a genomic region containing neighboring RNAP II (i.e. histone) and RNAP III (i.e. tRNA) genes, we observe specific binding of the associated Polymerase with shared transcription factors and chromatin modification patterns over all genes **(Figure 5C)**.

Although the presence of these histone modifications does not appear to distinguish between genes transcribed by different polymerases, we observed that both their position relative to the transcriptional start site (TSS) and their relative levels differ by polymerase: for RNAP I genes, these modifications localize prior to the TSS; for RNAP II, they flank the promoter and are enriched downstream of the TSS; and for RNAP III, they flank the gene body, localizing both upstream of the TSS and downstream of the transcriptional termination site **(Figure 5B, Figure S5D)**. In addition, the three RNAPs have different relative levels of these histone modifications near their respective gene promoters. For example, we found that RNAP I and II promoters display stronger H3K56Ac enrichment and RNAP I and III display stronger H3K4me2 enrichment relative to H3K4me3 **(Figure 5D)**. In this way, both quantitative combinations of histone modifications and their relative positions define distinct classes of promoters (**Figure 5E**).

Next, we considered whether other histone modifications may distinguish activity states of RNAP II promoters. To explore this, we quantified the levels of ten additional histone modifications at each genomic region containing H3K4me3 and grouped them using hierarchical clustering. We identified five sets of H3K4me3 enriched genomic regions; four are enriched with other histone modifications (sets 1-4) and one is not (set 5). The four co-occurring sets correspond to H3K4me3 along with: H3K27me3/H2AK119ub (set 1), H3K36me3/H3K79me (set 2), H3K9me3/H4K20me3 (set 3), or H3K4me1/H3K27ac (set 4) (**Figure 6A**). These sets correspond to promoters that exhibit distinct transcriptional activity (e.g., high versus low expression) (**Figure 6B**) and are enriched for distinct classes of RNAP II-transcribed genes, such as ribosomal protein and cell cycle genes (set 2), zinc finger protein (set 3) and long intergenic ncRNAs genes (sets 3 and 4) (**Figure 6C-G**)^13,86^. Consistent with the fact that H3K4me3 localization associates with functionally distinct classes of promoters, we found that different combinations of chromatin regulators that read, write, and erase H3K4me3 localize at distinct promoters (**Figure S4G**).

**Figure 6:**
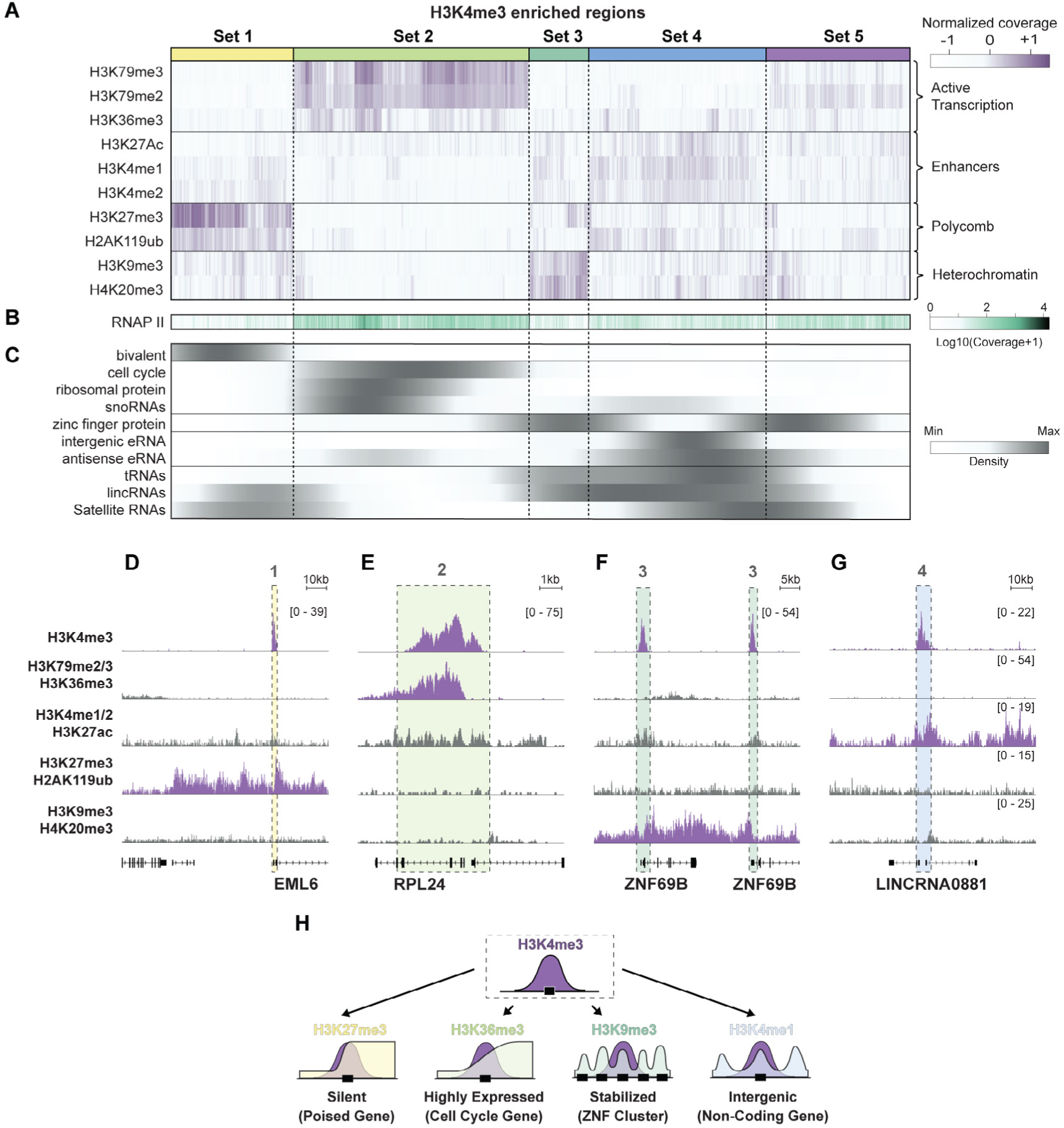
Combinations of histone modifications distinguish RNAP II promoter type, activity and potential. **(A)** Hierarchically clustered heatmap of coverage levels of 10 different histone modifications (y-axis) at individual H3K4me3 enriched genomic regions (x-axis). Five distinct sets of regions are indicated by colored bars along top-axis. **(B)** RNAP II coverage at H3K4me3-enriched regions, as sorted in (A). **(C)** Gene density of 10 different gene classes at H3K4me3 enriched regions, as sorted in (A). **(D)** Visualization of H3K4me3 and H3K27me3/H2AK119ub (associated with set 1) across the EML5 gene in K562. **(E)** Visualization of H3K4me3, H3K79me2/ H3K79me3/H3K36me3 colocalization (associated with set 2) across the ribosomal protein gene RPL24 in K562. **(F)** Visualization of H3K4me3 and H4K20me3/H3K9me3 colocalization (associated with set 3) across neighboring zinc finger genes in K562. **(G)** Visualization of H3K4me3 and H3K4me1/H3K4me2/H3K27Ac (associated with set 4) across the long intergenic noncoding RNA gene LNCRNA0881. For tracks in (D-G), the non-H3K4me3 tracks represent the sum of histone tracks associated with each set and are scaled to the maximum value across all panels. H3K4me3 tracks are scaled to the maximum for each panel. **(H)** Illustration summarizing the co-occurring promoter-associated histone modifications and their associated gene groups.

Taken together, these results demonstrate that combinations of histone modifications can distinguish promoter features including polymerase **(Figure 5E**), gene type, and activity level (**Figures 6H**).

### Enhancer type, activity and potential are defined by combinations of histone modifications

There are >40 different histone acetylation marks^87^, many of which have been associated with enhancers and active transcription. We mapped 15 of these in mESCs, including marks on all four core histones and histone variants, and observed that they co-localize at similar sites genome-wide (pearson r = 0.86-0.97)^88^ (**Figure S6**). We considered whether these strong correlations indicate that these marks are redundant or whether there is additional regulatory information encoded by the relative levels of each acetylation mark at specific genomic sites. To explore this, we used a matrix factorization algorithm to define five weighted combinations of acetylation marks at highly acetylated genomic regions (quantitative combinations C1-C5; see **Methods, Figure 7A-B, Supplemental Note 2, Figure S7)**. These quantitative combinations correspond to genomic regions that contain distinct transcription factor and chromatin regulator binding profiles **(Figure 7C-F, Figure S8)**.

**Figure 7:**
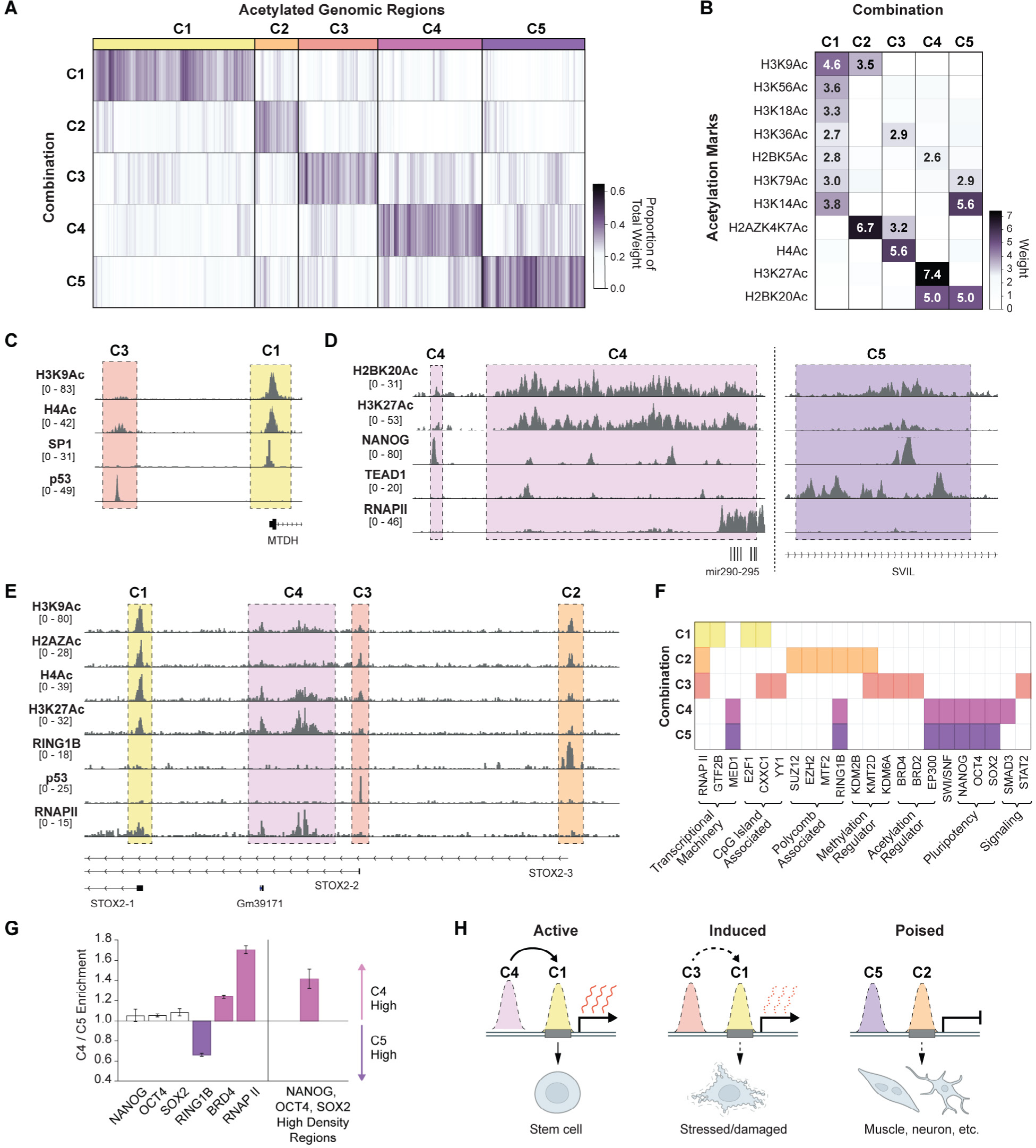
Combinations of histone modifications distinguish RNAP II promoter type, activity and potential. **(A)** The relative weights of five different combinations of histone acetylation marks (C1-C5, y-axis) for each acetylated genomic region (x-axis). Regions are grouped according to the combination that received the greatest weight, as indicated along top-axis. (**B)** The relative weights of each histone acetylation mark (y-axis) within each combination (x-axis). Only weights greater than 2.5 are labeled. **(C)** Visualization of H3K9ac and H4ac along with SP1 and P53 across a genomic region (mm10, chr15:34,065,000-34,086,000) containing enhancers assigned to the C1 (yellow) and C3 (red) state. **(D)** Visualization of H2BK20Ac and H3K27Ac along with Nanog, Tead1, and RNAP II across a genomic region (left: mm10, chr7:3,191,500-3,221,500, right: mm10, chr18:5,006,500-5,016,500) containing enhancers assigned to C4 (left) and a distinct region assigned to C5 (right). (Scale of the Nanog track is capped to the maximum of the left region; Tead1 data is from published ChIP-Seq data in fetal cardiomyocytes^96^). **(E)** Visualization of H3K9Ac, H2AZAc, H4Ac along with RING1B, P53, and RNAP II over a genomic region (mm10, chr8:47,272,800-47,427,000) containing enhancers assigned to states C1-C4. **(F)** DNA-associated proteins (x-axis, ordered by function) with significant binding at genomic regions defined by each combination (y-axis) are indicated in color (see **Methods**). **(G)** Enrichment bargraph of selected transcription-associated factors or regions with high density of pluripotency TFs (see **Methods**) in C4 vs C5 associated-regions. Error bars correspond to the enrichment range from bootstrap resampling. **(H)** Schematic of C1-C5 associated regions and their corresponding functions.

### Active promoter-proximal elements

The first group of regions (C1) is defined by H3K9Ac and several other H3 acetylation marks (H3K14ac, H3K18ac, H3K36ac, H3K56ac, and H3K79ac) (**Figure 7B)**. Genomic regions containing this signature tend to be localized near the promoter region of transcribed genes and are enriched for RNAP II, general TFs (e.g. TFIIB), and other CpG-island associated factors (e.g. E2F1, CXX1) along with their sequence motifs (e.g. ETS, SP and NRF families) **(Figure 7C,E-F)**.

### Poised promoter-proximal elements

The second group of regions (C2) contains high levels of H3K9Ac and acetylation of the histone variant H2AZ (H2AZAc) (**Figure 7B**). Genomic regions containing this signature tend to have lower levels of RNAP II (relative to C1) and are strongly enriched for polycomb (JARID2, SUZ12, RING1B) and other repressive chromatin regulators (KDM2B, HDAC2) **(Figure 7E-F)**.

#### Stress and signaling response elements

The third group of regions (C3) contains high levels of H2AZAc and H4Ac (**Figure 7B)**. Genomic regions containing this signature are also enriched for RNAP II but are bound by p53 and contain other stress response motifs (e.g BACH1, NRF2) or signaling response motifs (e.g. CRE) (**Figure 7C**, **Figure 7E-F**). Consistent with these observations, H2AZ has been proposed as a facilitator of inducible transcription (e.g. signaling pathway responses and p53 regulation)^89–92^. Yet, because H2AZ is also a component of C2, our results suggest that this association is not solely a property of H2AZAc but of this unique C3 signature.

### Active pluripotency distal regulatory elements

The fourth group of regions (C4) is defined by H2BK20Ac and H3K27Ac (**Figure 7B)**. Genomic regions containing this signature tend to be promoter-distal (**Figure S8B**) and are associated with actively transcribed embryonic and stem cell specific genes (**Figure 7D)**. These regions are enriched for binding of the pluripotency TFs, including Nanog, Oct4, and Sox2, as well as the P300 acetyltransferase and components of mediator **(Figure 7F)**.

### Poised differentiation distal regulatory elements

The fifth group of regions (C5) is defined by high-levels of H2BK20Ac (similar to C4) and H3K14Ac (distinct from C4) (**Figure 7B**). Interestingly, these regions displayed similar TF and CR occupancy (e.g. Oct4, Sox2, Nanog, P300 and mediator) to C4 regions **(Figure 7F-G)**. However, in contrast to C4 regions bound by these pluripotency factors, which correspond to enhancers of active genes involved in embryo and stem cell function, C5 regions bound by these factors correspond to enhancers of genes involved in post-embryonic development **(Figure S9)** and are enriched for sequence motifs of TFs involved in lineage specification and morphogenesis (e.g. TEAD family)^93^ **(Figure S8C)**. This suggests that C5 enhancers might be important in establishing the gene expression program needed upon differentiation (regulatory potential). Interestingly, we identified a third set of genomic regions that also contain a high-density of pluripotency TFs but lack the C4 or C5 acetylation signatures; these are associated with genes involved in later stages of organogenesis (e.g. kidney and sensory systems) (**Figure S9)**.

These analyses indicate that histone acetylation is not a redundant marker of enhancers, but that combinations of acetylation modifications can define unique classes of *cis* regulatory elements (promoter-proximal versus distal enhancers) that act in distinct ways (stimulus-responsive versus developmentally regulated) and that exhibit different activity (e.g. active gene expression versus poised for activation upon differentiation) (**Figure 7H**).

Overall, these observations highlight the importance of multi-component analyses and demonstrate why ChIP-DIP provides a powerful approach that will be critical for defining unique regulatory features within distinct cell states.

## DISCUSSION

We demonstrated that ChIP-DIP enables highly multiplexed mapping of hundreds of regulatory proteins to genomic DNA in a single experiment. Although the largest ChIP-DIP experiment in this study contained >225 distinct antibodies, this number was primarily limited by the availability of high-quality antibodies and we expect that ChIP-DIP could be performed using even larger pools of antibodies. Because this approach employs standard molecular biology techniques, we expect that it will be readily accessible to any lab without the need for specialized training or equipment. As such, we anticipate that ChIP-DIP will enable a fundamental shift from large consortia generating reference maps for a limited number of cell types to individual labs generating cell-type-specific maps within any specific experimental system of interest.

Given the important information encoded within quantitative combinations of histone modifications, chromatin regulators, and transcription factors, comprehensively mapping these factors across cell-types will be critical for studying gene regulation and for defining the putative effects of genetic variants associated with human disease. For instance, while specific regulatory states have been shown to be encoded by combinations of histone modifications (e.g. bivalent domains^4^), the details of such states (i.e. number and cell-type specificity) has remained largely unexplored. The large numbers of histone modifications and regulatory proteins have until now necessitated a tradeoff between mapping many marks in a few cell-types or a few marks in many cell-types. ChIP-DIP overcomes this by mapping hundreds of proteins in a single experiment. Moreover, due to the nature of split-pool barcoding used in ChIP-DIP and because there is negligible antibody-bead-chromatin dissociation during the procedure, ChIP-DIP can also be used to map protein binding within multiple samples simultaneously using distinct sets of antibody-oligo-labeled beads. While we did not directly emphasize this capability in this paper, several ChIP-DIP experiments described here were performed simultaneously by barcoding multiple sample conditions within the same experiment (e.g., crosslinking conditions, IP conditions). In addition to the increase in scale provided by mapping multiple proteins within a single sample, ChIP-DIP also reduces many sources of technical and biological variability associated with processing individual proteins and samples. This ability to measure regulatory proteins at scale, in multiple cell conditions, and with reduced sources of variability will enable large scale mapping of dynamic protein localization across distinct cell types and timepoints.

Beyond the multiplexing applications highlighted in this work, ChIP-DIP can be directly integrated into multiple existing split-pool approaches to create additional capabilities that are not currently possible. For example, we previously showed that we can map the 3D genome structure surrounding individual protein binding sites (SIP)^94^; integrating this approach with ChIP-DIP will enable mapping of the 3D structures that occur at hundreds of distinct protein binding sites simultaneously. Moreover, we previously developed a method to map 3D genome contacts within thousands of individual single cells using this same split-pool approach^36^. Integrating this single cell approach with ChIP-DIP will enable comprehensive mapping of hundreds of regulatory protein binding sites within many thousands of individual cells. Finally, we previously showed that split-and-pool barcoding can be used to simultaneously map the spatial proximity of DNA and RNA to measure ncRNA localization and the levels of nascent RNA transcription at individual DNA sites^95^. Integrating this approach with ChIP-DIP will enable the direct measurements of protein binding and transcriptional activity at individual genomic locations, providing a direct link between binding events and the associated transcription activity within the same cell. For these reasons, we expect that ChIP-DIP will represent a transformative new tool for dissecting gene regulation.

## ACKNOWLEDGEMENTS

We thank Shawna Hiley for editing, Inna-Marie Strazhnik for illustrations, Benjamin Yeh for generating the GitHub repository for the ChIP-DIP pipeline, and Olivia Ettlin for cell culture assistance. This work was funded by grants from the NIH (R01 HG012216; R01 DA053178; U01 DK127420), the Chan Zuckerberg Initiative Ben Barres Early Career Acceleration Award, and NIH UCLA-Caltech Medical Scientist Training Program (T32GM008042). Sequencing was performed in the Millard and Muriel Jacobs Genetics and genomics facility with support from Igor Antoshechkin and at the Broad Institute Genomics Platform.

## AUTHOR CONTRIBUTIONS

A.A.P., M.R.B., and M.G. conceived of ChIP-DIP; A.A.P and M.R.B. developed ChIP-DIP; A.A.P., I.N.G., and J.K.G. optimized ChIP-DIP; A.A.P and I.N.G generated data presented in this paper; I.N.G. developed the computational pipeline; I.N.G. performed data analysis and visualization; A.A.P., I.N.G., and M.G. generated figures and wrote the manuscript.

## SUPPLEMENTAL TABLES

**Table S1.** Antibody Pools and Read Counts for ChIP-DIP Experiments

**Table S2.** Correlations and Peak Overlap between ChIP-DIP and ENCODE

**Table S3.** Peak Overlap between ChIP-DIP Experiments with Antibody Pools of Different Sizes and Protein Composition

**Table S4.** Peak Overlap between ChIP-DIP Experiments with Various Amounts of Starting Cell Lysate

**Table S5.** Motif Analysis for Transcription Factors

**Table S6**. Antibody-ID Oligonucleotide Sequences

## METHODS

### Cell Lines, Cell Culture and Crosslinking

#### Cell lines used in this study

We used the following cell lines in this study: (i) Female mouse ES cells (pSM44 mES cell line) derived from a 129 × castaneous F1 mouse cross and (ii) K562, a female human lymphoblastic cell line (ATCC, Cat # CCL-243).

#### Cell Culture Conditions

i. pSM44 mES cells were grown at 37C under 7% CO2 on plates coated with 0.2% gelatin (Sigma, G1393-100ML) and 1.75 mg/mL laminin (Life Technologies Corporation, #23017015) in serum-free 2i/LIF media composed as follows: 1:1 mix of DMEM/F-12 (GIBCO) and Neurobasal (GIBCO) supplemented with 1x N2 (GIBCO), 0.5x B-27 (GIBCO 17504-044), 2 mg/mL bovine insulin (Sigma), 1.37 mg/mL progesterone (Sigma), 5 mg/mL BSA Fraction V (GIBCO), 0.1 mM 2-mercaptoethanol (Sigma), 5 ng/mL murine LIF (GlobalStem), 0.125 mM PD0325901 (SelleckChem) and 0.375 mM CHIR99021 (SelleckChem). 2i inhibitors were added fresh with each medium change. Fresh medium was replaced every 24-48 hours depending on culture density, and cells were passaged every 72 hours using 0.025% Trypsin (Life Technologies) supplemented with 1mM EDTA and chicken serum (1/100 diluted; Sigma), rinsing dissociated cells from the plates with DMEM/F12 containing 0.038% BSA Fraction V. K562 cells were purchased from ATCC and cultured in 1x DMEM (Life Technologies, # 11965118), 10% FBS (VWR, #97068-091), 100U/mL Penicillin/Streptomycin (Life Technologies, # 15140122), 1mM Sodium Pyruvate (Thermofisher, #11360070), 2mM L-Glutamine (Life Technologies # 25030081) at 37C and 5% CO in 15cm plates (USA Scientific # 5663-9160Q).

#### Cell Harvest

i. For harvesting pSM44 mESCs, cells were trypsinized by adding 5 mL of TVP (1 mM EDTA, 0.025% Trypsin, 1% Sigma Chicken Serum; pre-warmed at 37C) to each 15 cm plate and rocking gently for 3-4 min until cells start to detach. 25 mL of wash solution (DMEM/F-12 supplemented with 0.03% GIBCO BSA Fraction V, pre-warmed at 37C) was added to each plate to inactivate the trypsin. Detached cells were transferred into 50 mL conical tubes, pelleted at 330 g for 3 min, washed in 4 mL of 1X PBS per 10 million cells and then pelleted in 1X PBS in preparation for crosslinking. (ii) For harvesting K562s, the cell suspension was transferred to 50mL conical tubes, pelleted at 330 g for 3 min, washed with 4 mL of 1X PBS per 10 million cells and then pelleted in 1X PBS in preparation for crosslinking.

#### Cell Crosslinking

Cells were crosslinked in suspension with 1% Formaldehyde for 10 min at room temperature. For both cell lines, during crosslinking steps and subsequent washes, volumes were maintained at 4 mL of buffer or crosslinking solution per 10 million cells. Pelleted cells were resuspended in 1ml of 1X PBS per 10 million cells and pipetted to disrupt clumps of cells. Next, cells were crosslinked in suspension in a final volume of 4 mL of 1% formaldehyde (FA Ampules, Pierce 28906) diluted in 1X PBS per 10 million cells and rocked gently for 10 min at room temperature. Formaldehyde was immediately quenched with addition of 200 ml of 2.5 M glycine (Sigma G7403-250G) per 1 mL of 1% FA solution and incubated with gentle rocking for 5 min at room temperature. Cells were then washed three times with 0.5% BSA in 1X PBS that was kept at 4C. Finally, aliquots of 10 million cells were prepared in 1.7 mL tubes; these cell aliquots were pelleted, flash frozen in liquid nitrogen and stored in −80C until lysis.

### Nuclear Isolation and Chromatin Preparation

#### Nuclear Isolation

Crosslinked cell pellets (10 million cells) were lysed using the following nuclear isolation procedure: cells were incubated in 0.7 mL of Nuclear Isolation Buffer 1 (50 mM HEPES pH 7.4, 1 mM EDTA pH 8.0, 1 mM EGTA pH 8.0, 140 mM NaCl, 0.25% Triton-X, 0.5% NP-40, 10% Glycerol, 1X PIC) for 10 min on ice. Cells were pelleted at 850 g for 10 min at 4C. Supernatant was removed, 0.7 mL of Lysis Buffer 2 (50 mM HEPES pH 7.4, 1.5 mM EDTA, 1.5 mM EGTA, 200 mM NaCl, 1X PIC) was added and the sample was incubated for 10 min on ice. Nuclei were obtained after pelleting and supernatant was removed (as above). Then, 550 uL of Lysis Buffer 3 (50 mM HEPES pH 7.4, 1.5 mM EDTA, 1.5 mM EGTA, 100 mM NaCl, 0.1% sodium deoxycholate, 0.5% NLS, 1X PIC) was added and the sample was incubated for 10 min on ice prior to sonication.

#### Chromatin fragmentation and size analysis

Chromatin was fragmented via sonication of the nuclear pellet using a Branson needle-tip sonicator (3 mm diameter (1/8’’ Doublestep), Branson Ultrasonics 101-148-063) at 4C for a total of 2.5 min at 4-5 W (pulses of 0.7 s on, followed by 3.3 s off). To check the resulting DNA size distribution, a small aliquot of 20uL of sonicated lysate was then added to 80uL of Proteinase K buffer ((20 mM Tris pH 7.5, 100 mM NaCl, 10 mM EDTA, 10 mM EGTA, 0.5% Triton-X, 0.2% SDS) and reverse crosslinked at 80C for 30 minutes. DNA was isolated using Zymo IC DNA Clean and Concentrator columns and eluted in water. 10uL of purified DNA was then run for 10 minutes on a 1% e-gel (Invitrogen™ E-Gel™ EX Agarose Gels, 1%, Cat.No. G402021). Fragments were found to be 150-700 bp with an average size of roughly 350 bp. The remaining chromatin prep was stored at 4C overnight to be used for the immunoprecipitation the next day.

### ChIP-DIP: Bead Preparation

#### Antibody-ID oligo design

Antibody-ID oligos were designed and ordered from IDT (**Supplemental Table 6**). The sequence is as follows: /5phos/TGACTTGNNNNNNNNTATTATGGTAGATCGGAAGAGCGTCGTGTACACAGAGTC/3Bio/.

This corresponds to a sticky end that ligates Odd barcodes, UMI, antibody barcode, Illumina primer binding site (i5 primer binding site), spacer sequence. The oligo contains a 5’ phosphate to enable ligation and a 3’ biotin to enable coupling to beads.

#### Protein G Bead biotinylation

1 mL of Protein G Dynabeads (ThermoFisher, #10003D) were washed once with 1X PBSt (1X PBS + 0.1% Tween-20), separately keeping the original storage buffer, and resuspended in 1mL PBSt. Beads were then incubated with 20 μL of 5 mM EZ-Link Sulfo-NHS-Biotin (Thermo, #21217) on a HulaMixer for 30 minutes at room temperature. To quench the NHS reaction, beads were placed on a magnet, 500 μL of buffer was removed and replaced with 500 μL of 1M Tris pH 7.4 and beads were incubated on the HulaMixer for an additional 30 minutes at room temperature. Beads were then washed twice with 1 mL PBSt and resuspended in their original storage buffer until use.

#### Preparation of streptavidin-coupled oligonucleotides

Biotinylated antibody-ID oligonucleotides were coupled to streptavidin (BioLegend, #280302) in a 96-well PCR plate. In each well, 20 μL of 10 μM oligo was added to 75 μL 1X PBS and 5 μL 1 mg/mL streptavidin to make a 909 nM (calculated from the molarity of streptavidin molecules) stock. The 96-well plate was incubated with shaking at 1600 rpm on a ThermoMixer for 30 minutes at room temperature. Each well was diluted 1:4 in 1X PBS for a final concentration of 227 nM.

#### Preparation of oligonucleotide coupled Protein G beads

For each antibody in the experiment, 10uL of oligonucleotide-coupled Protein G beads were prepared. All biotinylated Protein G beads that would be needed for the entire experiment were first pooled into a tube, washed in 1mL of PBSt and resuspended in 200uL of 1x oligo binding buffer (0.5X PBST, 5 mM Tris pH 8.0, 0.5 mM EDTA, 1M NaCl) per 10uL of beads. 200 μL of bead suspension was aliquoted into individual wells of a deep well 96-well plate (Nunc 96-Well DeepWell Plates with Shared-Wall Technology, Thermo Scientific, Cat. No. 260251) and 14 μL of 5.675nM (1:40 from 227nM working stock made fresh) of streptavidin-coupled oligo was added to each well.

The 96-well plate was then sealed with a Nunc 96-well cap mat (Thermo Scientific, Cat. No. 276000) and shaken at 1200 rpm on a ThermoMixer for 30 minutes at room temperature. Beads in each well were washed twice with M2 buffer (20 mM Tris 7.5, 50 mM NaCl, 0.2% Triton X-100, 0.2% Na-Deoxycholate, 0.2% NP-40), twice with 1X PBSt, and finally resuspended in 200 μL of 1X PBSt.

#### Estimating number of oligos per bead

After oligo-coupling, a QC step was performed to estimate the number of oligos bound to each bead. 20% of a representative well for each row of the 96-well plate of oligo-coupled beads was isolated and the “Terminal” tag from split-and-pool barcoding was ligated onto the oligos in these aliquots. Then, half of the ligated product was PCR amplified for 10 cycles and purified using 1x SPRI beads. The purified DNA product was run on an Agilent Tapestation using a D1000 tape to estimate molarity and this molarity was used to calculate the total number of molecules post PCR. Using this post-PCR number and the number of cycles of PCR, the number of unique molecules pre-PCR was estimated^40^. Finally, the number of unique molecules was divided by the number of beads put into the PCR reaction (2.7*10^6 beads per 1uL of stock biotintylated protein G beads) to calculate the estimated oligos per bead.

#### Antibody Coupling

2.5 μg of each antibody was added to each well of the 96-well plate containing oligonucleotide labeled beads resuspended in 1X PBSt. The plate was incubated on a ThermoMixer overnight at 4C with 30 seconds of shaking at 1200 RPM every 15 minutes. The following morning, beads were washed twice with 1X PBSt (Sigma, #B4639-5G), resuspended in 200 μL of 1x PBSt + 4mM biotin + 2.5ug Human IgG Fc and left shaking at 1200 rpm for 15 minutes at room temperature to quench free Protein G or streptavidin binding sites.

#### Preparation of bead pool

All wells containing oligo labeled, antibody coupled beads were washed 2X with 200uL 1X PBSt + 2 mM biotin, taking care to remove all supernatant after the final wash. Afterwards, one of two protocols were followed for bead pooling: 1) Equal bead pooling - Beads were pooled using equal amounts of prepared beads for each antibody (10uL of Protein G beads per antibody). 2) Titrated bead pooling - Beads were pooled using unequal amounts of prepared beads for each antibody. The relative number of beads for each antibody was determined based on the chromatin pull-down efficiency (chromatin reads per bead) measured in QC experiments. Fewer beads were used for antibodies with higher pull-down efficiencies and greater beads were used for antibodies with lower pull-down efficiencies or negative controls. This strategy was intended to generate a more uniform distribution for the number of chromatin reads assigned to each antibody in the final experiment.

### ChIP-DIP: Immunoprecipitation, Split-and-pool and Library Preparation

#### Pooled immunoprecipitation

Fragmented lysate was diluted with PBSt +10mM biotin + 1x PIC + 2.5ug of human IgG Fc per 10ul of beads. The pool of labeled beads was added to the lysate and rotated on a HulaMixer for 1 hour at room temperature. Beads were washed 2X with 1mL IP Wash Buffer I (20mM TrispH8.0, 0.05% SDS, 1% Triton X 100, 2mM EDTA, 150mM NaCl in water), 2X with 1mL of IP Wash Buffer II (20mM TrispH8.0, 0.05% SDS, 1% Triton X 100, 2mM EDTA, 500mM NaCl in water) and 2X with 1mL of M2 buffer (20mM Tris pH7.5, 0.2% Triton X100, 0.2% NP-40, 0.2% DOC, and 50mM NaCl).

### Chromatin End Repair and dA-tailing

To blunt end and phosphorylate double stranded DNA, the NEB End Repair Modu.le (E6050L; containing T4 DNA Polymerase and T4 PNK) was used. Beads were incubated in 1X NEBNext End Repair Enzyme cocktail + 1X NEBNext End Repair Reaction Buffer + 4mM biotin + 1ug human IgG Fc per 10uL beads at 20C for 15 minutes. The reaction was quenched with 3X volume of PBSt + 100uM EDTA and beads were washed 2X with 1mL PBSt. Next, to dA-tail DNA, the NEBNext dA-tailing Module (Klenow fragment (50 −30 exo-, NEBNext dA-tailing Module, E6053L) was used. Beads were incubated in 1X NEBNext dA-tailing Reaction Buffer + 1X Klenow Fragment (exo-) + 4mM biotin + 1ug human IgG Fc per 10uL beads at 37C for 15 minutes. The reaction was quenched with 3X volume of PBSt + 100uM EDTA and beads were washed 2X with 1mL PBSt.

#### Split-and-pool barcoding

Split-and-pool barcoding was performed as previously described^38,40^ with modifications. Specifically, beads were first split-and-pool ligated by DPM to attach a common sticky end to all DNA molecules. Then, beads were split-and-pool ligated for ≥ 6 rounds with sets of “Odd,” “Even,” and “Terminal” tags. The number of barcoding rounds and number of tags used for each round was determined based on the number of beads that needed to be resolved. These parameters were selected to ensure that virtually all barcode clusters (>95%) represented molecules belonging to unique, individual beads. In most cases, 6 rounds of barcoding with 24-36 tags per round were performed. Each individual tag sequence was used in only a single round of barcoding. All split-and-pool ligation steps were performed for 5 minutes at room temperature and supplemented with 2mM biotin and 5.4uM ProteinG. After split-and-pool barcoding was complete, beads were resuspended in 1mL of MyRNK buffer [20 mM Tris pH 7.5, 100 mM NaCl, 10 mM EDTA, 10 mM EGTA, 0.5% Triton-X, 0.2% SDS]. Aliquots of various sizes (0.05% to 4% of total beads) were prepared, ensuring that the number of beads within each aliquot was resolvable by the number of possible unique split-and-pool barcodes. Each aliquot was then digested with 8ul of Proteinase K (NEB) for 2 hrs at 55C, 1200RPM shaking and reverse crosslinking at 65C, 1600 RPM shaking overnight.

#### Library Preparation

DNA from each reverse crosslinked aliquot was isolated with a Zymo IC column using a 6X volume of the DNA binding buffer (Zymo Cat. No. D4014) and eluted in 21ul of H20. Libraries were amplified for 9-12 cycles using Q5 Hot-Start Mastermix (NEB Cat No M0294L) and primers that added the full Illumina adaptor sequences. The following PCR mixture was used: 21uL DNA in H2O, 2uL of i5 primer (12.5uM), 2uL of i7 primer (12.5uM), 25uL 2X Q5 MM. After amplification, libraries were cleaned with 1.2x SPRI (Bulldog Bio CNGS500) and eluted in 20uL. Prior to sequencing, libraries were gel purified to remove unused primers using a 2% agarose gel [Invitrogen Cat No. G401002].

### Sequencing

Sequencing was performed on Illumina NovaSeq S4 (300 cycle) and NextSeq (200 cycle or 300 cycle) paired-end runs, Read lengths were asymmetrical in order to capture the full split-and-pool barcode sequence on read 2 (R2) and the chromatin sequence on read 1 (R1). For 300 cycle kit – 100 cycles for R1, 200 cycles for R2; For 200 cycle kit – 50 cycles for R1 and 150 cycles for R2.

For each experiment, multiple different libraries were generated and sequenced. Each library corresponds to a distinct aliquot which is amplified with a unique pair of primers, providing an additional round of barcoding.

### Data Processing Pipeline

#### Read Processing

Paired-end sequencing reads were trimmed with Trim Galore! V0.6.2 (https://www.bioinformatics.babraham.ac.uk/projects/trim_galore/) to remove adaptor sequences and quality assessed with FastQC v0.11.8. Split-and-pool barcodes were identified from Read 2 using Barcode ID v1.2.0 (https://github.com/GuttmanLab/chipdip-pipeline). Reads missing split-and-pool tags or with tags in the incorrect position given the split-and-pool round they correspond to were discarded. Subsequently, reads were split into two files, one for antibody ID reads and one for DNA reads, based on the presence of “BPM” (bead tag) or “DPM” (DNA tag), respectively, on Read 1.

For DNA reads, the DPM sequence was trimmed from Read 1 using Cutadapt v3.4^99^. The remaining sequence was aligned to mm10 or hg38 using Bowtie2 (v2.3.5)^100^ with default parameters. Only primary alignments with a mapq score of 20 or greater were kept for further analysis. Finally, reads were masked using the repeat genome obtained from ENCODE^101^.

For antibody ID reads, the BPM sequence, which contains the antibody-ID information, was trimmed from Read 1 using Cutadapt v3.4 and the UMI extracted from the remaining sequence.

MultiQC v1.6^102^ was used to aggregate metrics from all steps.

#### Cluster Generation

A “cluster file” was generated by aggregating all reads (ie. aligned, masked DNA reads and antibody ID reads) that share the same split-and-pool barcode sequence. During this step, reads in each cluster were deduplicated by alignment position for DNA reads or UMI for antibody ID reads.

#### Antibody ID Oligo Movement Quality Control Check

To assess the frequency of antibody ID oligo movement between beads, the proportion of antibody ID reads corresponding to the maximum representation in each cluster was calculated. Only clusters with >1 antibody ID read were considered. For each experiment, these values were plotted as an empirical cumulative distribution function (ECDF) using the python plotting package seaborn^103^.

#### Cluster Filtering and Assignment

Individual clusters in the “cluster file” were assigned to a specific antibody based on antibody ID reads within the cluster. First, clusters in the “cluster file” without antibody ID reads or clusters with >10,000 genomic DNA reads (which likely represent undersonicated material or clumps of beads) were filtered out. Next, each remaining cluster was assigned to the antibody ID that had maximum representation within the cluster if 1) there were greater than two unique reads corresponding to the antibody ID and 2) the antibody ID represented >80% of all antibody ID reads within the cluster. These criteria were selected empirically to ensure high confidence assignments of antibody IDs to each cluster. Clusters that did not meet these criteria were removed from further analysis.

#### Antibody-specific protein maps

Genomic DNA alignments were split into separate bam files such that each file corresponded to all alignments associated with an individual antibody based on the antibody ID assignments within each cluster. DNA reads from clusters that did not have antibody ID reads, were too large or could not be uniquely assigned to a single antibody ID were filtered out. DNA reads were deduplicated such that only one read per alignment position per cluster was retained.

### Visualization and Peak Calling

#### Bigwig Generation

Bigwigs were generated from each antibody-specific BAM file using the ‘bamCoverage’ function from deeptools v3.1.3^104^ and were visualized with IGV^105^. Track visualizations are scaled to the maximum over the region and scales indicate reads per bin, unless indicated otherwise.

#### Background Normalization

Because beads from many antibodies are processed together, ChIP-DIP has sources of potential background that are distinct from a traditional ChIP-Seq experiment. In any ChIP experiment, the antibody used will immunoprecipitate its specific protein (and the associated chromatin) but will also non-specifically purify other proteins (and their associated chromatin) at some lower frequency. This non-specific chromatin (background) is generally proportional to the overall distribution of genomic DNA present in the starting material (“input”). In ChIP-DIP, the same is true during the IP; any given antibody will preferentially capture its specific protein but will, at some lower frequency, non-specifically capture other proteins. However, because ChIP-DIP entails purification with many antibodies, the source of proteins and chromatin for this non-specific binding is no longer the entire cellular input but rather the material present within the pooled IP (e. g. the proteins and chromatin that were pulled down by the pool of antibodies). Indeed, we observed that some antibodies displayed background signal that was distinct from the input library. For example, antibodies targeting CTCF displayed higher background over promoter regions, likely reflecting the presence of various promoter-enriched histone modifications present in the same experimental pool. To account for this in our analysis, we used the pool of all genomic DNA reads captured in a ChIP-DIP experiment as the background control. We found that normalization using this empirically-defined background led to a more conservative enrichment calculation for ChIP-DIP data. For example, in the CTCF example noted above, this normalization approach successfully removed the background promoter-associated signal while retaining signal at known CTCF binding sites.

Specifically, a background model was generated for each individual antibody using the total pool of assigned sequencing reads. The background for an antibody contained all reads except those assigned to it, or other antibodies targeting the same or related proteins. For example, for an antibody targeting RNAPII-NTD, reads from all antibodies targeting RNAP II were excluded from this background set. To calculate a scaling factor for this background: 1) the total experiment coverage was calculated in 10kB bins, 2) the high coverage bins (80%) were selected, 3) a per-bin enrichment quotient of the target compared to the background coverage was calculated, 4) a kernel density plot of the enrichment quotient was generated, 5) a threshold was calculated based on the position of the smallest peak and 6) the ratio of total coverage in target versus background bins below the threshold was determined. The goal of this procedure was to locate regions that represented background noise in the target and calculate the target-to-background ratio using only those regions. The kernel density plot was frequently bimodal or with a long tail, with the higher peak or tail representing signal bins and the lower peak representing background bins. Background normalized peaks were called using the scaled background as a substitute for input. Background normalized bigwigs were generated using the ‘bamCompare’ function from deeptools v3.1.3 by subtracting the scaled background and, subsequently, removing negative value bins.

#### Peak Calling

Peaks were called using the HOMER v4.11^106^ program ‘findPeaks’ on tag directories generated for target datasets using ‘-style histone’ for histone modification targets and ‘-style factor’ for other targets. Background normalized peaks were generated using the scaled background distribution (described above). Specific parameter settings, such as ‘-minDist’ (distance between adjacent peaks), ‘-size’ (width of peaks) or filtering thresholds were tuned according to the nature of the target. For instance, peaks for focal histone modification H3K4me3 were generated using ‘-F 2-P 0.001 -L 0’ while enriched regions for broad histone modification H3K36me3 were calculated using ‘-F 2 -P 0.001 -L 0 -size 1000 -minDist 7500 -region’.

#### Motif Prediction

Transcription factor motifs were predicted using the HOMER program ‘findMotifsGenome’ on peaks generated using HOMER, as described above. For all transcription factors, motifs were generated using the settings ‘-s 200 -mask −l 10’ and results are reported in **Supplemental Table 5**. For individual examples in **Figure 4** with, longer motifs were also predicted.

### Ribosomal DNA Alignments

To analyze reads aligning to genomic DNA encoding ribosomal RNA (rDNA), we aligned reads directly to an rDNA reference. We generated a modified reference of the mouse rDNA sequence (NCBI Genbank BK000964.3)^107^. Because the original mouse BK000964.3 sequence begins with the TSS and ends with the Pol I promoter, we transposed a portion at the end of the rDNA reference to the beginning, as previously described^108^, to enable a continuous visualization of the promoter-TSS region. Specifically, the rDNA sequence was cut at the 36,000 nt position and the sequence downstream of the cut site were moved upstream of the TSS, such that the resulting rDNA sequence begins with ∼10kb of IGS, then the promoters and then transcribed regions. Processing steps prior to sequence alignment followed the standard ChIP-DIP pipeline. After barcode identification, DNA sequence was aligned to the custom rDNA genome using Bowtie2 (v2.3.5) with default parameters. Only primary alignments with a mapq score of 20 or greater were kept for final analysis. The subsequent cluster generation and read assignment steps followed the standard ChIP-DIP pipeline.

#### ChIP-DIP Experiments

We performed 9 ChIP-DIP experiments in this paper, each of which, along with the associated antibodies, proteins, and statistics are described in **Supplemental Table 1**. Briefly, these experiments were:

*1. <u>Chromatin Movement Experiment</u>*: A quality control human and mouse mixing experiment used to quantify possible chromatin movement during the procedure.
*2. <u>Antibody Movement Experiment</u>:* A quality control human and mouse mixing experiment used to quantify possible antibody movement during the procedure.
*3. <u>K562 10 Antibody Pool Experiment:</u>* An initial data-generation experiment performed in human K562 to measure a small number of well-defined targets.
*4. <u>K562 50 Antibody Pool Experiment:</u>* A data-generation experiment performed in human K562 measuring 50 antibodies.
*5. <u>K562 52 Antibody Pool Experiment:</u>* A data-generation experiment performed in human K562 measuring 52 antibodies.
*6. <u>K562 35 Antibody Pool Experiment:</u>* A data-generation experiment in human K562 measuring 35 antibodies as a function of different cell input amounts.
*7. <u>mESC 228 Antibody Pool Experiment:</u>* An antibody-screening experiment performed in mouse ES cells using 228 antibodies to identify good antibodies and the antibody amounts required for deeper characterization in subsequent ChIP-DIP experiments.
*8. <u>mESC 67 Antibody Pool Experiment:</u>* A data-generation experiment performed in mouse ES cells measuring 67 antibodies.
*9. <u>mESC 165 Antibody Pool Experiment:</u>* A data-generation experiment performed in mouse ES cells measuring 165 antibodies.

All ChIP-DIP experiments were performed using the same general protocol with the following experiment-specific modifications:

## 1. Chromatin Movement Experiment

To test whether chromatin dissociates during the ChIP-DIP procedure and binds to other beads, we designed a human-mouse mixing experiment. Cell lysate from 20M mESC cells, cell lysate from 10M K562 cells and two sets of antibody-coupled, oligonucleotide-labeled beads were prepared according to standard protocol. Prior to IP, lysate yields were quantified using TapeStation and equal amounts of mouse and human chromatin preparations were used for the subsequent, separate IPs. One set of antibody-ID labeled beads was used for the human IP and the other set of antibody-ID labeled beads was used for the mouse IP. After IP, the two species-specific IPs were mixed and split into three conditions using different quenchers: (i) 10% BSA quencher, (ii) 1X Blocking Buffer quencher and (iii) No quencher. For the 10% BSA quencher condition, end-repair, dA tailing and DPM reactions were performed in buffer supplemented with 10% BSA. For the 1X blocking buffer quencher condition, end-repair, dA tailing and DPM reactions were performed in buffer supplemented with 1X protein blocking buffer (Abcam ab126587). The three conditions were combined for split-and-pool barcoding.

For alignment of human-mouse mixing experiments, DNA reads were aligned to a custom combination genome including both mm10 and hg38 genomes using Bowtie2 (v2.3.5) with default parameters. Only primary alignments with a mapq score of 20 or greater were kept for further analysis. Reads were then masked using a merged version of mm10 and hg38 blacklist regions defined by ENCODE. Reads were then uniquely assigned to human beads (beads present only in the human IP condition) or mouse beads (beads present only in the mouse IP condition) using the standard assignment pipeline. Total reads aligned to mm10 or hg38 for each bead set were quantified and the relative proportions were plotted as a bar graph.

### 1. 2. Antibody Movement Experiment

To test whether antibodies dissociate from their labeled beads during the ChIP-DIP procedure and bind to other beads, we designed an experiment that involved the addition of labeled antibody-free beads at various steps. Following a similar set-up as the chromatin mixing experiment, cell lysate from 20M mESC cells, cell lysate from 10M K562 cells and two sets of antibody-coupled, oligonucleotide-labeled beads were prepared using the standard protocol. One set of beads was used for the human IP and the other set of beads was used for the mouse IP. After IP, half of each species-specific IP was mixed together, and half was left separate. For this mixed condition only, oligonucleotide-labeled beads without a coupled antibody were added prior to the end repair and the dA-tailing reactions. These empty beads were added to capture antibodies that dissociated from other IP’d beads. Finally, the three conditions (mouse only, human only, mixed) were ligated with unique sets of DPM adaptors and combined for split-and-pool barcoding. To calculate the frequency of antibody movement, total reads and total beads assigned to human CTCF beads, human IgG beads, empty beads added prior to end repair and empty beads added prior to dA tailing were quantified. Reads per bead for each group were normalized to the mean value for human CTCF beads. These normalized values were plotted as a bar graph with 99% CI using the python plotting package seaborn.

### 1. 3. K562 10 Antibody Pool Experiment

We performed an initial small scale proof-of-concept (POC) experiment in K562 using 10 different antibodies. The POC experiment was performed using lysate from 50M K562 cells per IP. Standard protocol with equal bead pooling was used with the exception of IP conditions. Two identical sets of antibody coupled beads were prepared using different biotinylated oligonucleotides; one set was used for an overnight immunoprecipitation at 4C and one set was used for 1-hr immunoprecipitation at room temperature. DNA processing steps and DPM ligation reactions were performed separately for the two IP conditions and then the two samples were pooled for the remaining rounds of split-and-pool barcoding. See **Supplemental Table 1** for full list of antibodies under the “K562 10 Antibody Pool” tab. For data processing, the standard pipeline generated individual clusters corresponding to antibody-IP condition pairs and individual bam files for each target in each IP condition. Data from both IP conditions were merged for each target, resulting in a single file per antibody.

### 1. 4. K562 50 Antibody Pool Experiment

The K562 50 Antibody Pool Experiment was performed using lysate from 50M K562 cells. The standard protocol with equal bead pooling was used. See **Supplemental Table 1** for full list of antibodies under the “K562 50 Antibody Pool” tab.

### 1. 5. K562 52 Antibody Pool Experiment

The K562 52 Antibody Pool Experiment was performed using lysate from K562 cells. To test the efficiency of different crosslinking strategies, two parallel IPs were performed using the same pool of prepared beads. One IP utilized 60M K562 cells crosslinked with 1% FA and the other IP utilized 60M K562 cells crosslinked with 1% FA + DSG. Cells for the 1% FA condition were prepared as described above. Cells for the 1% FA + DSG condition were prepared as follows: After harvest and pelleting, K562 cells were crosslinked in 4 mL of 2 mM disuccinimidyl glutarate (DSG, Pierce) dissolved in 1X PBS per 10 million cells for 45 minutes at room temperature. Cells were then pelleted, washed with 1X PBS and crosslinked with 1% FA, as described above.

For antibody ID oligonucleotide-labeling of beads, beads were labeled in two sequential rounds. First, beads were labeled according to the standard protocol. Then, beads were labeled again using another 2.5uL of 5.67nM streptavidin-coupled oligo in 200uL of 1x oligo binding buffer for 30 minutes at room temperature. During the first round of labeling, all wells received a unique streptavidin-coupled oligonucleotide. During the second round of labeling, most wells received the same streptavidin-coupled oligonucleotide as the first round, with the exception of eleven wells. Eleven pairs of wells received the same streptavidin-coupled oligonucleotide in the second round – one well of each pair was labeled with the same oligonucleotide in both rounds while the other well was labeled with different oligonucleotides. The result was that most beads were labeled with a single, unique oligonucleotide label, eleven beads were labeled with a pair of oligonucleotide labels, and eleven beads were labeled with a single oligonucleotide label that can also be found on other beads. This labeling strategy was designed to test combinatorial labeling of beads. After antibody coupling, beads were pooled in equal amounts and half of the bead pool was used for IP of each crosslinking condition. Following IP, each condition was processed separately and DPM-ligated with unique, condition-identifying sets of adaptors. Conditions were kept separate for the first round of split-and-pool barcoding and then combined for the remaining rounds of split-and-pool. See **Supplemental Table 1** for full list of antibodies under the “K562 52 Antibody Pool” tab.

Sequenced data was processed using the standard ChIP-DIP pipeline up until the clustering assignment step. To account for the dual oligo labeling of selected antibodies, prior to assignment of unique antibodies to each cluster, clusters with multiple labels (clusters containing both oligo types from a known co-occurring pair) were isolated and antibody-ID oligos in these clusters corresponded to the second round of labeling were reassigned to the matched antibody-ID oligo from the first round of labeling. The result is that all antibodies now corresponded to a unique antibody-ID oligo; for the eleven combinatorial pairs, this is the first round of labeling. Afterwards, the remaining steps in the standard ChIP-DIP pipeline (cluster assignment, etc) were performed as described above.

### 1. 6. K562 35 Antibody Pool Experiment for input cell number titration

One of the major challenges with mapping DNA binding proteins in primary cell types, disease models, and other rare cell populations is the large number of cells required for traditional ChIP-Seq experiments. Because ChIP-DIP enables simultaneous mapping of many proteins within the same experiment, we reasoned that it may dramatically reduce the total number of cells required in two ways: (i) the number of cells required to map any individual protein is instead distributed across all protein targets in a pool, and (ii) the total chromatin purified from multiple proteins may enable purification of lower DNA concentrations associated with a single/low abundance proteins that might otherwise be lost due to experimental handling.

The K562 35 Antibody Pool Experiment was designed to measure the amount of cell input material required for ChIP-DIP. To do this, we performed a series of ChIP-DIP experiments using the same antibody pool and differing amounts of cell lysate. Specifically, we crosslinked >100M cells in a single batch and then performed four ChIP-DIP experiments from this same crosslinked lysate. This experiment involved four separate ChIP-DIP experiments, performed in pairs of two. For the first pair, the 45M and 5M conditions, a 50M cell aliquot was lysed and sonicated and then split into 45M and 5M cell equivalents of lysate. For the second pair, the 500K and 50K conditions, a 1M cell aliquot was lysed and sonicated and then split into 500k and 50k cell equivalents of lysate. Each pair of experiments used a single preparation of antibody-coupled, antibody ID oligonucleotide labeled beads that was split in half. See **Supplemental Table 1** for full list of antibodies under the “K562 35 Antibody Pool” tab.

Read coverage profiles of four targets – H3K4me3, H3K27me3, CTCF and RNAP II - were compared. For both RNAP II and CTCF, two different antibodies were included (RNAP II: CST 91151 and 14958S; CTCF: CST 3418S and ABCAM ab128873). Coverage of normalized bigwig files across the set of all peak regions from the 10 Antibody Pool experiment, the same set of regions used for the pool size comparison correlations, was calculated using the ‘multiBigwigSummary’ function of the python package deeptools v.3.1.3. Pearson correlation coefficients for all pairs were calculated using the ‘plotCorrelation’ function of deeptools v.3.1.3 and the plotted as a heatmap, manually ordering the rows/columns from lowest to highest amount of input lysate for each target.

Peak overlaps were calculated for each antibody between pairs of experiments as the (number of peaks in experiment 1 intersecting peaks in experiment 2) / (total number of peaks in experiment 1). The number of intersecting peaks were calculated using the bedtools v2.29.2 and results are reported in **Supplemental Table 4**.

### 1. 7. mESC 228 Antibody Pool Screen

The mESC 228 Antibody Pool experiment was performed using 40M mESC cells. The standard protocol was used with the following modifications. Because the number of antibodies exceeded the number of unique antibody-ID oligonucleotides, three plates of antibody-coupled, oligonucleotide-labeled beads were prepared separately, the beads from each plate were pooled into three separate antibody-bead pools, and each pool was used to IP one third (∼13M cells) of the prepared cell lysate. Post IP, each sample was processed separately until split-pool. During split-pool barcoding, a unique set of “ODD” barcodes was ligated to each sample during the first round and then all samples were pooled for rounds 2-6.

Sequencing data was processed through the standard pipeline, using a concatenated string of three antibody names (ie. DNMT3B-CST_POLR3E-Bethyl_H3K36Ac-CST), one name for the antibody corresponding to each sample, to match individual antibody-ID sequences during barcode identification. After cluster generation and prior to cluster assignment, each antibody-ID read was assigned to only one antibody based on its first-round ‘ODD’ split-and-pool tag. Cluster assignment then proceeded using the standard pipeline. Chromatin reads were assigned to each antibody based on cluster assignments and total chromatin for each antibody was quantified (see **Supplemental Table 1**, “mESC 228 antibody pool” tab for counts). These relative chromatin yields per antibody were used to inform the ideal amount of antibody needed and we used these to titrate the amount of beads for each antibody pooled together in ChIP-DIP experiments 8 and 9 below. This experiment was sequenced at low depth to allow for rapid and low-cost antibody screening.

### 1. 8. mESC 67 Antibody Pool Experiment

The mESC 67 Antibody Pool experiment was performed using lysate from 80M mESC cells. The standard protocol with titrated bead pooling was used. This experiment contained 67 different antibodies. See **Supplemental Table 1** for full list of antibodies under the “mESC 67 Antibody Pool” tab.

### 1. 9. mESC 165 Antibody Pool Experiment

The mESC 165 Antibody pool experiment was performed using lysate from 60M mESC cells. Similar to the mESC 228 Antibody Pool experiment, because the number of antibodies exceeded the number of unique antibody-ID oligonucleotides, a multi-plate strategy was used. Specifically, two plates of antibody-coupled, oligonucleotide-labeled beads were prepared separately, pooled using the titrated bead pooling strategy and used to IP half of the prepared cell lysate. After IP, the two samples were processed separately up until the third round of split-and-pool barcoding and then combined for the remaining rounds of split-and-pool. This experiment contained 165 different antibodies. See **Supplemental Table 1** for full list of antibodies under the “mESC 165 Antibody Pool” tab. Sequencing data was processed through the standard pipeline, using a concatenated string of antibody names (ie. LSD1-CST_SAP30-Bethyl) to match individual antibody-ID sequences during barcode identification. After cluster generation and prior to cluster assignment, each read antibody-ID read was assigned to only one antibody based on its first-round split-and-pool tag. Cluster assignment and BAM file generation then proceeded using the standard pipeline.

### Protein Target Classification

Antibody targets were assigned to one of five categories: histone modification (HM), transcription factor (TF), chromatin regulator (CR), RNA polymerase (RNAP) and other DNA associated protein. Transcription factors were defined as proteins with a DNA-binding domain and were manually subclassified into constitutive, stimulus response or cell type specific/developmental manually curated based on functional descriptions from GeneCards. Chromatin regulators contained proteins or members of complexes that read, write, or erase histone modifications or DNA methylation. Proteins that were part of chromatin regulator complexes and contained a DNA binding domain were considered part of the chromatin regulatory category. Proteins involved in chromatin remodeling (e. g. BRG1) or other structural proteins that interact with chromatin (e. g. LaminA) were also considered chromatin regulators. Dual function proteins (e. g. transcription factors with intrinsic acetyltransferase capabilities) were assigned to a single category (e. g. transcription factor) but were included in chromatin regulator schematics. Other DNA associated proteins included a mixture of targets, such as RNAP elongation factors (e. g. ELL), RNA binding proteins (e. g. NONO) and antibodies that detected DNA methylation.

### Comparison to ENCODE data

ChIP-DIP comparisons to ENCODE-generated ChIP-Seq data in **Figure 1** were performed using the 10 pool experiment in K562. Visual comparisons were performed using IGV and the raw ENCODE datasets: ENCFF656DMV (H3K4me3), ENCFF785OCU (POLR2A), ENCFF800GVR (CTCF) and ENCFF508LLH (H3K27me3). Genome-wide coverage comparisons were calculated across all RefSeq TSS for H3K4me3 and POLR2A or across 10kB bins for CTCF and H3K27me3. Calculations were performed using the ‘multiBamSummary’ function of the python package deeptools v3.1.3 and plotted as 2-D kernel density plots using the python library seaborn.

Systematic comparisons between ChIP-DIP K562 datasets (10 Antibody Pool, 50 Antibody Pool, 52 Antibody Pool and 35 Antibody Pool) and ENCODE-generated ChIP-Seq were performed for all targets for which ENCODE datasets were available. Genome-wide coverage comparisons were calculated at 1000bp using the ‘multiBigwigSummary’ function of the python package deeptools v3.1.3 and Pearson correlation coefficients are reported in **Supplemental Table 2**. The number of overlapping peaks between ChIP-DIP and ENCODE datasets were calculated using as described above. Both the fraction of ENCODE peaks detected by ChIP-DIP and the fraction of ChIP-DIP peaks detected by ENCODE are reported in **Supplemental Table 2**.

Accession numbers used for coverage comparisons include: ENCFF121RHF, ENCFF508LLH, ENCFF035SOZ, ENCFF272JVI, ENCFF816ECC, ENCFF880HKV, ENCFF352HXD, ENCFF446FUS, ENCFF656DMV, ENCFF465UWC, ENCFF149MXA, ENCFF155UQU, ENCFF187HIQ, ENCFF702HIC, ENCFF800GVR, ENCFF982AFE, ENCFF178ARN, ENCFF096FWU, ENCFF844WTT, ENCFF801TEZ, ENCFF174BEG, ENCFF014HSG, ENCFF656FDC, ENCFF457PGP, ENCFF108EMO, ENCFF617EYG, ENCFF986KSB, ENCFF572IXE, ENCFF893LSE, ENCFF785OCU, ENCFF816VGU, ENCFF179XDZ Accession numbers used for peak comparisons include: ENCFF863MYY, ENCRR698RKX, ENCRR736TRL, ENCFF222OPH, ENCFF731WDM, ENCFF479VOH, ENCFF514SHW, ENCFF643TQX, ENCFF586UHP, ENCFF456SZC, ENCFF108JNI, ENCFF632MQY, ENCFF924JXT, ENCFF834RTA, ENCFF660HBV, ENCFF567BOX, ENCFF563CWK, ENCFF394JNY, ENCFF307DMW, ENCFF181AVH, ENCFF169ZQQ, ENCFF040TWS, ENCFF990EZP, ENCFF755KWM, ENCFF269JZL, ENCFF106GOL, ENCFF827OVS, ENCFF789NDS, ENCFF567HEH

### Pool Size Comparison Analysis

To measure the influence of the number of antibodies contained within an individual pool, read coverage profiles of four targets – H3K4me3, H3K27me3, CTCF and RNAP II – generated in four different ChIP-DIP experiments in K562 cells were compared. ChIP-DIP experiments included the10 Antibody Pool, the 45M condition from the 35 Antibody Pool, the 50 Antibody Pool and the 52 Antibody Pool in K562. For both RNAP II and CTCF, two different antibodies were included (RNAP II: CST 91151 and 14958S; CTCF: CST 3418S and ABCAM ab128873).

Coverage of normalized bigwig files across the set of all peak regions from the 10 Antibody Pool experiment was calculated using the ‘multiBigwigSummary’ function of the python package deeptools v.3.1.3. Pearson correlation coefficients for all pairs were calculated using the ‘plotCorrelation’ function of deeptools v.3.1.3 and plotted as a heatmap, manually ordering the rows/columns from smallest to largest pool size for each target.

Peak overlaps were calculated for each target between experiments of different pool sizes as described above and are reported in **Supplemental Table 3**.

### Histone Modification Diversity Analysis

#### Chromatin-State

Genome-wide coverage for 10kb windows for 12 histone marks (H3K27me3, H2AK119ub, H3K9me3, H4K20me3 and H3K9me3 from the 5M condition in the 35 Antibody Pool Experiment in K562; H3K79me2, H3K79me1, H3K4me3, H3K4me2, H3K4me1, H3K9Ac and H3K27Ac from the histone panel in K562) was calculated using the ‘multiBamCoverage’ function from deeptools v3.1.3. These values were standardized for each mark by transforming into z-score values. The UMAP reduction was generated using the UMAP^109^ python package and parameters n_components=2 and n_neighbors=3.

#### Polycomb-Associated Histone Modifications

Validation of polycomb-associated histone modifications used the 5M condition in the K562 35 Antibody Pool Experiment. H3K27me3 and H2AK119ub bam alignment files were converted into binary signal files using the ‘BinarizeBam’ script from the ChromHMM^110^ package with standard settings. The number of bins with only H2AK119ub signal or with both H2AK119ub and H3K27me3 signal were computed and plotted as a pie chart.

#### Heterochromatin-Associated Histone Modifications

Validation of heterochromatin-associated histone modifications used the 5M condition in the K562 35 Antibody Pool Experiment. Read coverage of H3K9me3, H4K20me3 and H3 were computed over annotation groups (ZNFs, LTRs, LINES, SINES, TSS+/-2kb) using the ‘depth’ function from samtools v1.9^111^. An enrichment score was calculated by normalizing for feature and target abundance. Specifically, let a = total base pairs within an annotation group, b = effective genome size, c = read coverage of a target over the annotation group and d = total reads of the target. The enrichment score would be (c/d) / (a/b).

#### Promoter-Associated Histone Modifications

Validation of promoter-associated histone modifications used the ChIP-DIP histone dataset in mESC. Promoter coverage correlations were calculated across promoters from EPDNew^112^, a database of non-redundant eukaryotic RNAP II promoters, +/-500bp using the ‘multiBamSummary’ and ‘plotCorrelations’ functions of the python package deeptools v.3.1.3.

#### Gene Body-Associated Histone Modifications

Validation of gene body-associated histone modifications used the 5M condition in the K562 35 Antibody Pool Experiment and the K562 50 Antibody Pool Experiment. Coverage metaplots over the gene bodies of all protein coding genes from GENCODE v38 basic annotation were calculated using ‘computeMatrix’ function of the python package deeptools v.3.1.3 and normalized to the maximum and minimum for each target.

#### Enhancer-Associated Histone Modifications

Validation of enhancer-associated histone modifications used the 5M condition in the K562 35 Antibody Pool Experiment and the K562 50 Antibody Pool Experiment. H3K4me1 peaks were assigned to three categories (promoter, gene or intergenic) based on overlap with H3K4me3 (promoter), H3K79me1 (gene) or H3K36me3 (gene). These categories were further sub-divided based on the co-occurrence of H3K27Ac peaks. The proportion of peaks in each category was computed and plotted as a pie chart.

### Chromatin Regulator Diversity Analysis

#### Polycomb-Associated Chromatin Regulators

Validation of polycomb-associated chromatin regulators used the K562 50 Antibody Pool Experiment. Metaplots respective to RING1B peak sites were calculated using ‘computeMatrix’ function of the python package deeptools v.3.1.3 with the following settings: ‘reference-point -bs 10000 -a 500000 -b 500000’. The resulting read coverage profiles were normalized to the maximum and minimum for each target and plotted as a heatmap.

#### Heterochromatin-Associated Chromatin Regulators

Validation of heterochromatin-associated chromatin regulators used the K562 50 Antibody Pool Experiment. Genome-wide coverage for 10kB windows and Pearson correlation coefficients were calculated using the ‘multiBigwigSummary’ function and ‘plotCorrelation’ function, respectively, of the python package deeptools v3.1.3.

#### H3K4me3-Associated Chromatin Regulators

Analysis of H3K4me3-associated chromatin regulator used the mESC 165 Antibody Pool Experiments. Binding profiles of JARID1A, RBBP5 and PHF8 were measured +/-1kB around the TSS of all representative promoters from EPDNew and were clustered using k-means clustering with k=4 by the ‘plotCoverage’ function of the python package deeptools v.3.1.3. H3K4me3 binding profiles from the mESC 67 Antibody Pool Experiment were measured over the same four promoter groups.

### Polymerase Diversity Analysis

#### RNAP I, II and III Comparison

Validation of the various RNA polymerases used the mESC 165 Antibody Pool Experiment. First, read coverage within a +/-100bp window surrounding the promoters/TSS of various gene groups were calculated. For tRNAs, the TSS of repeatmasker^113^ tRNAs were used. For snRNAs, the TSS of repeatmasker snRNAs (excluding U6 which is transcribed by RNAP III) were used. For mRNAs, EPDNew TSS annotations were used. For rDNA, the spacer promoter was used. Next, for each polymerase, coverage was normalized to the total reads aligned with any gene group. Finally, an enrichment score of the relative coverage compared to IgG was calculated and plotted as a bar graph.

#### RNAP II Phosphorylation State Comparison

Validation of the various RNA polymerases used the K562 52 Antibody Pool Experiment. Metaplots over the gene bodies of all protein coding genes from GENCODE v38 basic annotation were calculated using ‘computeMatrix’ function of the python package deeptools v.3.1.3.

### Histone Combinatorial Analyses

#### Polymerase-Associated Histone Profiles

For RNAP I, track coverage profiles of various histone modifications 1.5kB upstream to 0.5kB downstream of the spacer promoter were visualized using IGV. For RNAP II, metaplots of coverage profiles for various histone modifications were generated around active and inactive RNAP II promoters using the deeptools v.3.1.3 ‘computeMatrix’ (reference-point -a 1000 -b 1000) and ‘plotProfile’ functions. Promoters were defined as the TSS of all representative promoters from EPDNew and were grouped into active or inactive based on the read coverage of RNAP II in the surrounding +/-1kB window.

For RNAP III, metaplots of coverage profiles for various histone modifications were generated around active and inactive tRNA genes using the deeptools v.3.1.3 ‘computeMatrix’ (scale-regions -a 1000 -b 1000 -m 75 -bs 25) and ‘plotProfile’ functions. tRNA genes were grouped into active or inactive based on the read coverage of RNAP III.

For comparison of relative histone levels, total coverage for each histone mark was calculated in the −1.5kB to +0.5kB window surround the spacer promoter for rDNA, −0.5kB to +0.5kB window around active RNAP II promoters and −0.5kB to +0.5kB window around active RNAP III tRNA gene promoters. To account for differences in window size, the coverage of H3K56Ac and H3K4me2 was normalized to the level of H3K4me3. The density profiles of these ratios were plotted using the seaborn ‘jointplot’ function with the following kde parameters: “common_norm=False, thresh=0.2, log_scale=True, levels=10, cut=True”. For comparison to RNAP I, the total sum ratios (e. g. total H3K4me2 coverage across all active RNAP II promoter intervals divided by total H3K4me3 coverage across all active RNAP II promoter intervals) were also calculated and plotted for RNAP II and RNAP III.

#### H3K4me3 Enriched Regions Clustering

Combinatorial histone modification analysis for H3K4me3 regions used the 5M condition of the K562 35 Antibody Pool Experiment. Read coverage of ten histone targets (H3K79me3, H3K79me2, H3K36me3, H3K4me1, H3K4me2, H3K27Ac, H3K27me3, H2AK119ub, H3K9me3 and H4K20me3) was calculated over all H3K4me3 peak regions using the ‘multicov’ function of bedtools^114^. The resulting region vs histone data matrix (A) was normalized using log normalization^115^: 1) The log of the data matrix was computed *L=log log (A)*. 2) The column mean (*<u>L</u>_i_*), row mean (*<u>L</u>_j_*), and overall mean (*<u>L</u>*) of the log matrix were computed. 3) All individual cells of the final matrix were computed according to K_ij_=L_ij_ - *<u>L</u>_i_ - <u>L</u>_j_ + <u>L</u>*. This method of normalization is intended to capture the “extra” coverage of histone modification *j* in region *i* that is not explained simply by the overall difference between region *i* and other regions or between histone modification *j* and other histone modifications. Instead, it is special to the combination of region *i* (a region with H3K4me3 enrichment) and coverage of histone modification *j*. The regions of the normalized data matrix were clustered using cluster.hierarchy.linkage function from scipy v.1.6.2^116^ with a Euclidean distance metric and complete linkage method. The clustered matrix was visualized using the ‘clustermap’ function of python package seaborn.

Gene annotation of H3K4me3 regions was performed using the ‘annotatePeaks.pl’ function from HOMER v4.11. ZNF genes, RP genes, and lincRNA genes were defined as regions whose annotation gene description contained the terms ‘zinc finger protein’, ‘ribosomal protein’ and ‘long intergenic’, respectively, and had the nearest TSS within 2000bp. snoRNA genes were defined as all regions whose annotation gene type was snoRNA. Satellite RNA genes were defined as regions whose detailed annotation contained the term ‘Satellite’. tRNA genes were defined as all regions that intersected with tRNA gene bodies or upstream by 500bp of the tRNA TSS from repeat masker. Cell cycle genes were defined as regions whose gene annotation belonged to the Kegg Cell Cycle Pathway^117^. Bivalent genes were defined as regions whose gene annotation belong to those identified by Court and Arnaud in human H1 cells^118^. Enhancer RNA regions (both antisense and intergenic) were defined as regions that intersected those identified by Lidschreiber et al.^119^ and had the nearest TSS greater than 2000bp away. To visualize enrichments of gene annotations in sets and subsets of the hierarchically clustered heatmap, the kernel density estimate (KDE) was calculated for each annotation group based on their clustering-defined order.

RNAP II levels of individual H3K4me3 regions were measured as the summed coverage over each region from four antibodies targeting RNAP II (RNAP II, RNAP II NTD, RNAP II Ser5, RNAP II Ser2) from the K562 52 Antibody Pool Experiment. Transcriptional levels for sets and subsets of H3K4me3 regions were compared using violin plots generated by the python plotting package seaborn. P-values for comparison of transcriptional levels within subsets of H3K4me3-enriched regions were calculated using the kolmogorov smirnov test from scipy.stats.

### ChromHMM Model of Acetylation

The ChromHMM genome segmentation model was built using 15 different histone acetylation modifications measured in the mESC 67 Antibody Pool Experiment. Bam files were binarized using the BinarizeBam function from ChromHMM with a poisson threshold of 0.000001 and other default parameters. The signal threshold was increased from default to remove spurious noise. State models with 5-20 states were built using the LearnModel function with default parameters. States were manually reordered and grouped based on transition probabilities between states. 19 states were selected for the final model to retain state 17, a state with a distinctive enrichment and transition profile.

### Non-Negative Matrix Factorization of Acetylated Regions

Non-negative matrix factorization analysis utilized the histone acetylation mark data from the mESC 67 Antibody Pool Experiment. NMF is a matrix factorization technique to reduce dimensionality and explain the observed data using a limited number of combinatorial components^115^. NMF decomposes the original data matrix (dimensions: N x M) into a basis matrix (dimensions: N x k) and a mixture coefficient matrix (dimensions: k x M). In this case, N represents genomic regions of interest, M represents individual histone acetylation marks and k represents the number of combinatorial histone acetylation states. High coverage regions were defined using the results of the ChromHMM Model. Specifically, the 200bp genomic bins corresponding to states with enrichment of multiple histone acetylation marks (states 1,2,3,4,6,9,10,11,12,15,16) were merged to form high coverage regions. Then, to reduce the number of fragmented or spurious regions, bins with 400 base pairs (2 genomic windows) were merged and regions with size less than 400 base pairs (2 genomic windows) were filtered out. A initial normalized data matrix (N x M) was generated by computing the coverage of each histone modification over each region and normalizing for region size and histone abundance. Specifically, to account for differences in region size between regions, the total reads per region was scaled by region size and, to account for differences in total measured histone abundance between marks, sigmoidal scaling was used^120,121^. NMF was then performed using ‘Nimfa’^122^, a python library for nonnegative matrix factorization, with the nndsvd initialization method. The rank *k* was selected empirically, taking into account the biological assignability of the resulting states, the complexity of the model and the stability of the factorization (the number of iterations the algorithm required to coverage).

After factorization, the resulting basis matrix (N x k) contained the coefficient of each combination *i* for each genomic region. A sorted heatmap of the basis matrix was generated by grouping the regions according to the combination that contributed the greatest coefficient for each region. For visualization, this heatmap was normalized by dividing the coefficients for each region by the total coefficient sum of the region.

To profile and assign a biological interpretation to individual combinations, each region was assigned to the combination with the maximum coefficient. Identification of transcription factors with significant binding overlap to regions assigned to a single combination was performed using the Cistrome Data Browser, an interactive database of public ChIPseq^123^. For each combination, the top 100 scores were filtered for targets with at least 2 hits in any cell type. Motif enrichment was calculated using the HOMER function ‘findMotifs’ on all genomic regions assigned to each combination. For comparison of enrichment levels in C4 versus C5, enrichments were calculated using bedgraphs from the mESC 165 Antibody Pool Experiment and the ChromHMM program ‘OverlapEnrichment’ (java-jar ChromHMM.jar OverlapEnrichment-binres 1-signal). Interval bars for these enrichments were generated by bootstrap resampling; enrichments were recalculated for 200 independent draws of 75% of the regions assigned to C4 or C5.

### High Density Regions of NANOG-OCT4-SOX2

High density regions of pluripotency associated transcription factors were calculated using the NANOG, OCT4 and SOX2 data from the mESC 165 Antibody Pool Experiment. Specifically, high-density and low-density regions were defined using the super-enhancer setting of the ‘callPeaks’ function from HOMER on the merged tag directors of the three transcription factors. To remove nonspecific background peaks, the merged tag directories of the background models for these three factors was used as input. Briefly, the super enhancer setting with default parameters first identifies peaks, then stiches together individual peaks that are within 12.5kb of each other, calculates a ‘super enhancer score’ for each region based on input-normalized read coverage, generates a ‘super enhancer plot’ (regions sorted by score vs number of regions) and identifies the regions where the slope of the plot is greater than 1. These regions are labeled as putative ‘super enhancers’ while all remaining regions are labeled as ‘typical enhancers’. We consider the ‘super enhancer’ regions as high-density regions (HDR) and the ‘typical enhancer’ regions as low-density regions (LDR).

TF and CR enrichments over HDRs versus LDRs were calculated using the ‘computeMatrix’ function with scale-regions setting from deeptools v.3.1.3. To account for the differences in typical region size between LDRs and HDRs, which tended to be much larger, the -m parameter was set to approximately the median region size for each group.

GO terms associated with the intersection of HDRs, LDRs and NMF-based acetylation combinations were calculated using the GO analysis function of ‘annotatePeaks’ from HOMER. To limit the number of terms under consideration, only terms assigned to the biological process category that received a cutoff p<0.001 were used. Terms were then manually grouped into larger categories (e. g. developmental, metabolic). Enrichment scores were calculated by normalizing for the total number of possible unique terms assigned the category and the total number of terms assigned to the intersection group.

### Statistics

Pearson correlation coefficients for coverage comparisons versus ENCODE were calculated using pearsonr function of scipy.stats library^116^. Pearson correlation coefficients for heatmaps were generated using the ‘plotCorrelation’ function from deeptools v.3.1.3^104^.

---

**Supplemental Figure 1:**
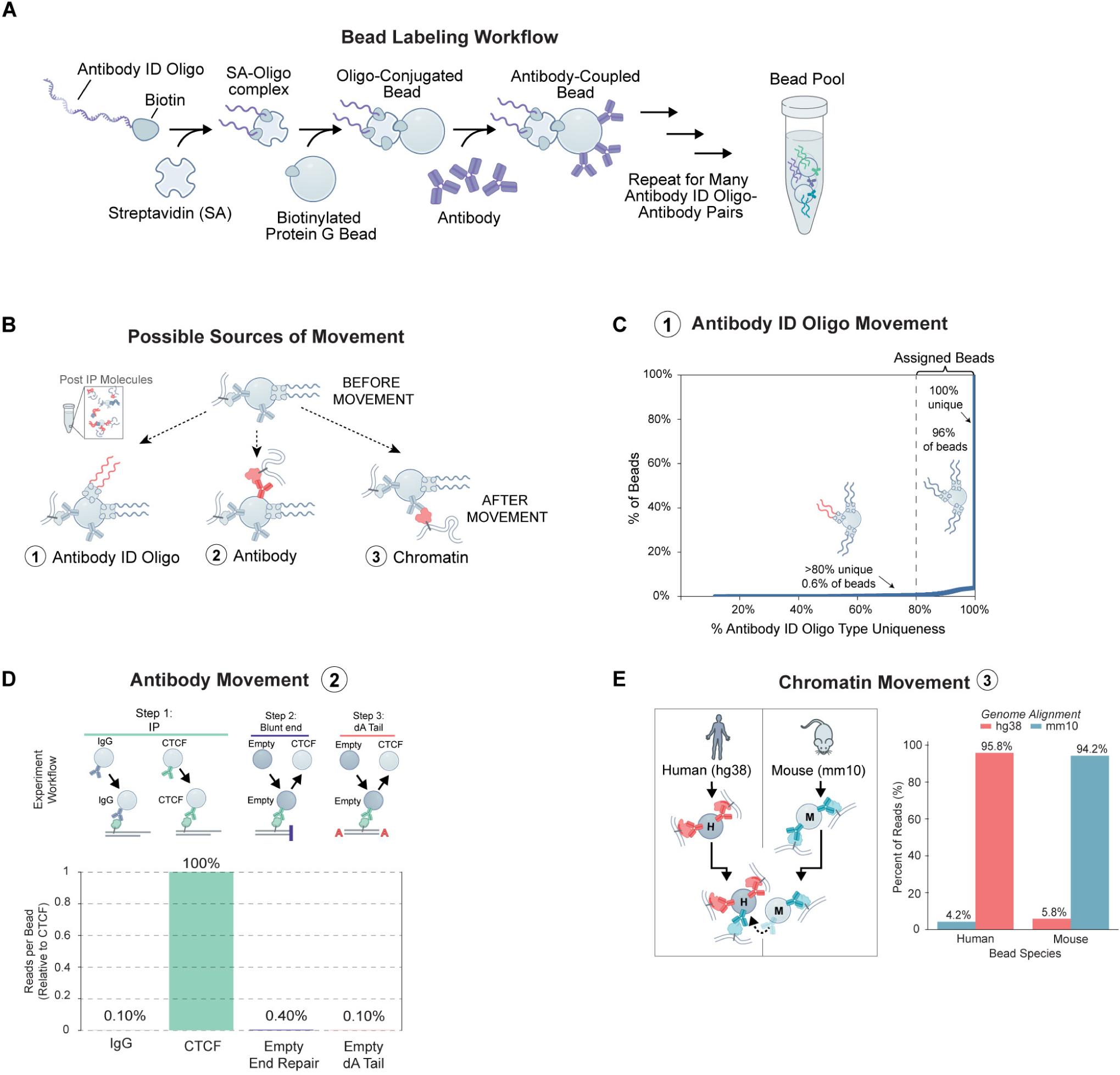
Potential sources of mixing in ChIP-DIP, related to Figure 1. **(A)** Schematic of labeling strategy to generate Protein G beads coupled with a unique antibody-identifying oligonucleotide and a matched antibody. Protein G beads are covalently modified with a biotin; oligonucleotides containing a 3’ biotin are conjugated to streptavidin; oligo-streptavidin complexes are mixed with biotinylated protein G beads and protein G beads are mixed with antibodies. This process is repeated for each unique oligonucleotide-antibody pair and pooled together. **(B)** Schematic of three potential sources of mixing during ChIP-DIP. **(C)** Cumulative distribution plot representing the uniqueness of antibody-ID oligos type (x-axis) within individual clusters. **(D)** Schematic of experimental design to test for antibody movement between beads and quantification of relative reads per bead assigned to true targets (CTCF) or empty beads added during experimental processing steps. **(E)** Schematic of human-mouse experimental design to test for chromatin movement and quantification of species-specific reads assigned to human or mouse beads.

**Supplemental Figure 2:**
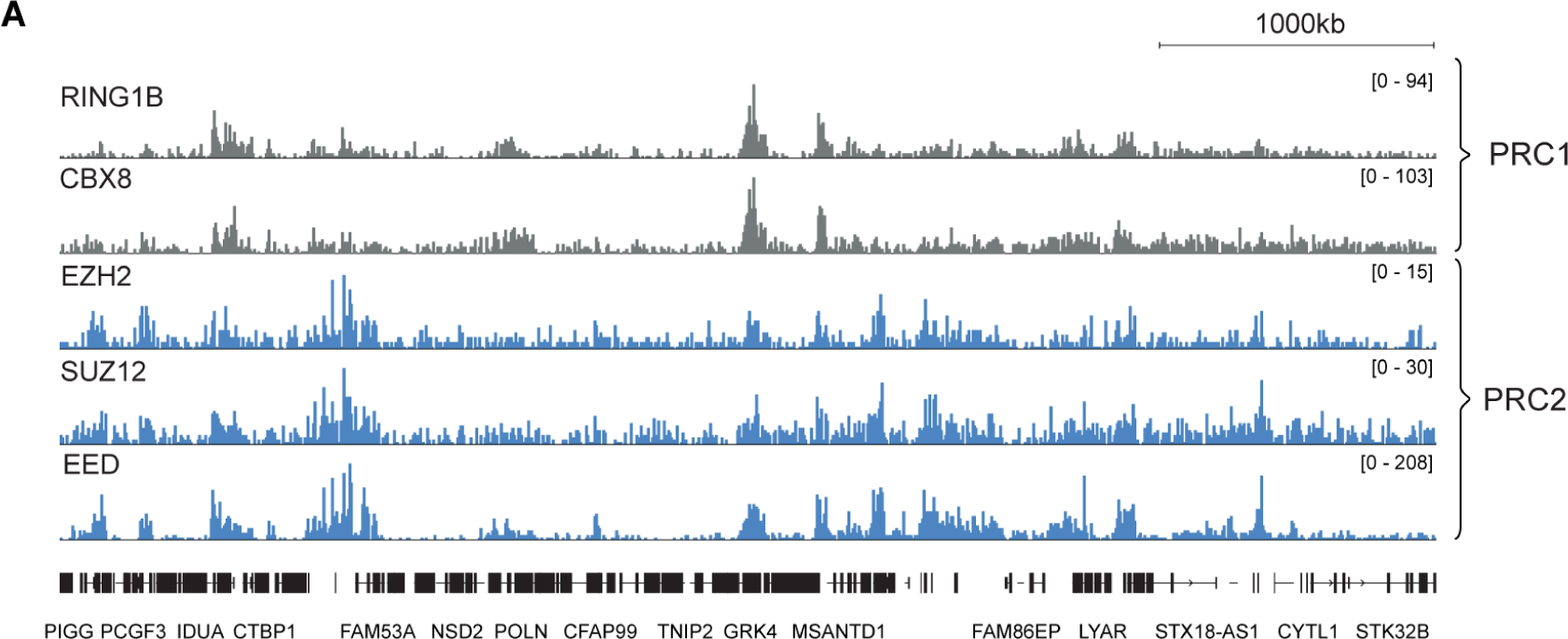
Simultaneous mapping of multiple components of a single protein complex using ChIP-DIP, related to Figure 2. **(A)** Visualization of various components of the PRC1 (RING1B, CBX8) and PRC2 (EZH2, SUZ12, EED) complexes that were mapped within the same ChIP-DIP pool (K562 52 Antibody Pool) along a genomic region (hg38, chr4:500,000-5,500,000).

**Supplemental Figure 3:**
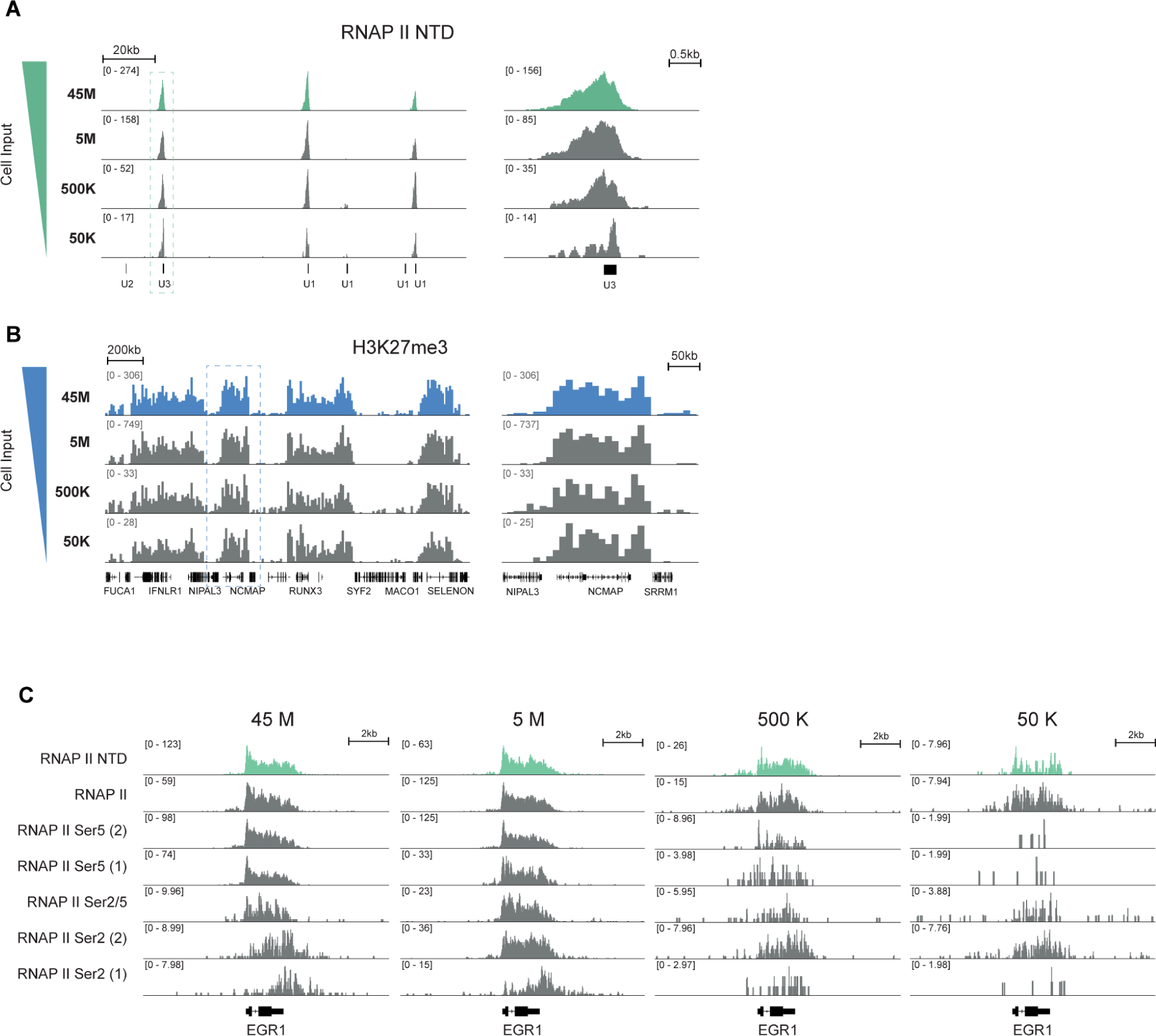
Comparison of protein localization across different amounts of cell lysate, related to Figure 2. **(A)** Comparison of RNAP II NTD localization across a snRNA gene cluster (hg38, chr17:58,620,000-58,689,000) when generated using various amounts of input K562 cell lysate. **(B)** Comparison of H3K27me3 localization across a genomic region (hg38, chr1:23,850,000-25,850,000) generated using various amounts of input K562 cell lysate. **(C)** Comparison of various isoforms of RNAP II at the EGR1 locus (hg38, chr5:138,455,000-138,480,000) generated using various amounts of input K562 cell lysate.

**Supplemental Figure 4:**
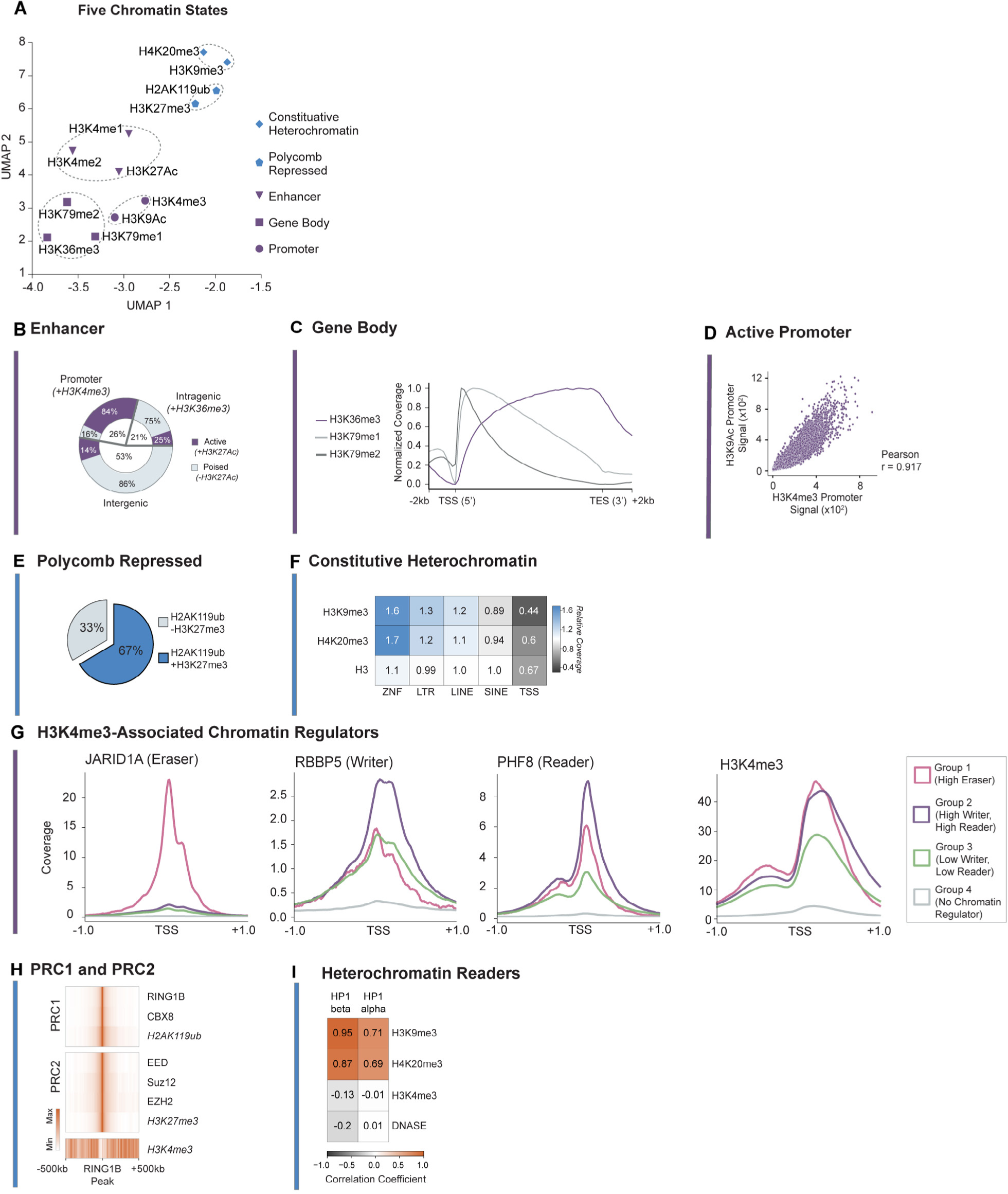
Histone modifications associated with five chromatin states, related to Figure 3. **(A)** UMAP embedding of 12 histone modifications measured in K562 correspond to five chromatin states. **(B)** Pie chart showing proportion of H3K4me1 peaks in K562 at various genomic position categories - promoters (co-localizing H3K4me3), intragenic (co-localizing H3K36me3) or intergenic - in K562. Proportion of peak regions overlapping with H3K27Ac within each genomic position category are shown in purple. **(C)** Metaplot of signal distribution of H3K36me3, H3K79me1 and H3K79me2 across the gene body of protein coding genes in K562. **(D)** Correlation scatterplot of H3K9Ac and H3K4me3 signals at promoter sites in mESC. **(E)** Pie chart showing overlap of H2AK119ub and H3K27me3 sites in K562. **(F)** Enrichment heatmap of H3K9me3 and H4K20me3 at various associated (ZNF genes, LTRs, LINES) and unassociated (SINES, TSS) genomic elements in K562. H3 is shown as reference. **(G)** Metaplots of read coverage for three H3K4me3-associated chromatin regulators (JARID1A, RBBP5, PHF8) and H3K4me3 at four promoter groups in mESC. Promoter groups were identified using k-means clustering of CR signal (see **Methods**). **(H)** Metaplot showing colocalization of multiple PRC1 and PRC2 members and their respective histone modifications at RING1B sites in K562. **(I)** Genome-wide correlation matrix of multiple HP1 proteins versus heterochromatin and euchromatin markers in K562.

**Supplemental Figure 5:**
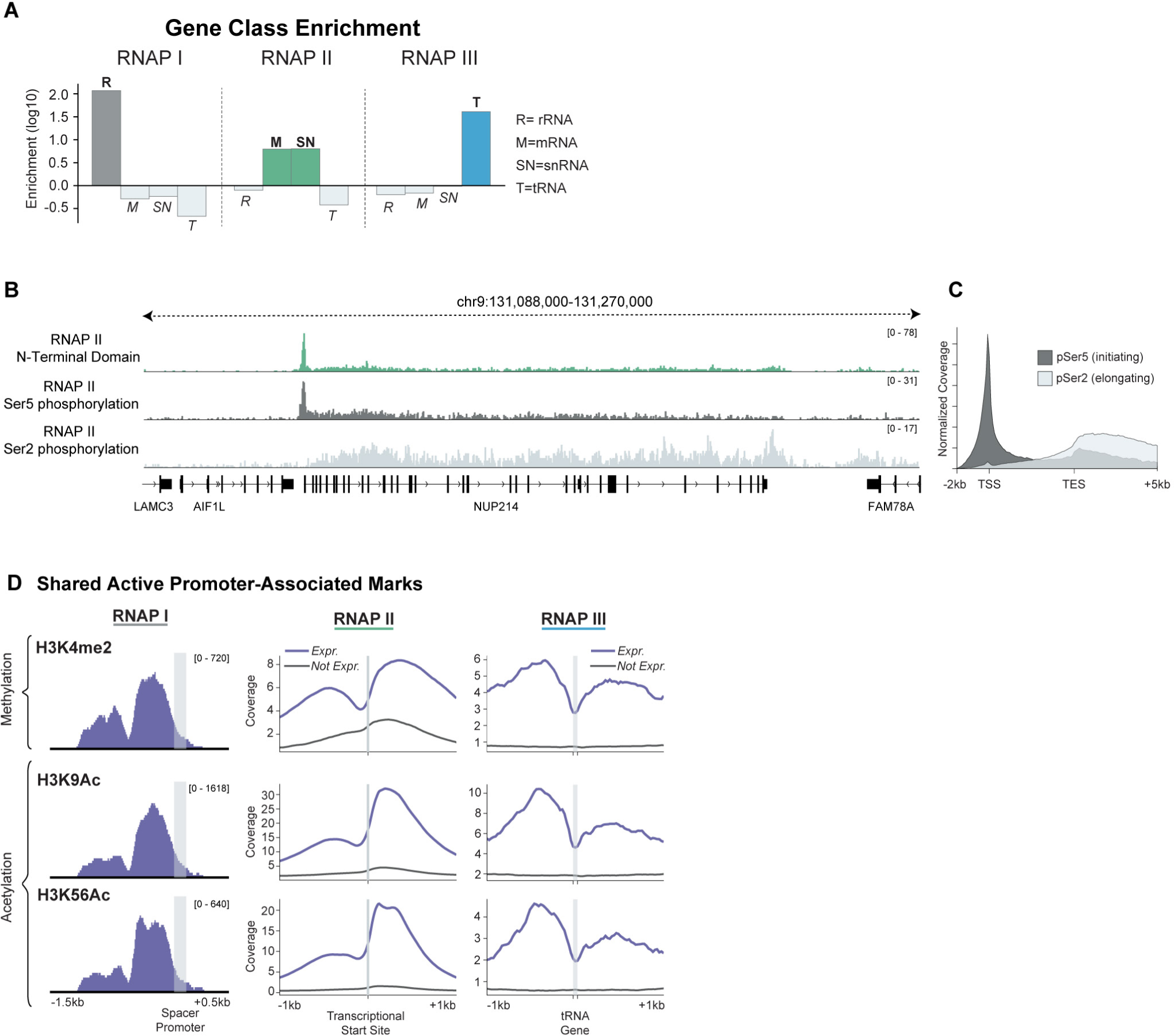
Chromatin states corresponding to distinct RNA polymerases and isoforms, related to Figure 5. **(A)** Bar graph showing enrichment of gene class coverage (rRNA, mRNA, snRNA or tRNA) for RNAP I, II and III in mESC. For each RNAP, the bar of its associated class (or classes) is highlighted. **(B)** Visualization of RNAP II phosphorylation isoforms across the NUP214 gene in K562. **(C)** Metaplot of signal distribution of RNAP II phosphorylation isoforms across the gene body of protein coding genes in K562. **(D)** Comparison of histone profiles for H3K4me2, H3K9Ac and H3K56Ac at the promoters of RNAP I, II and III, similar to Figure 5B. (Left) Histone modification over the RNAP I-transcribed rDNA spacer promoter. (Middle/Right) Metaplot of histone profiles at active (blue) and inactive (gray) promoters for RNAP II (middle) and RNAP III (right).

**Supplemental Figure 6:**
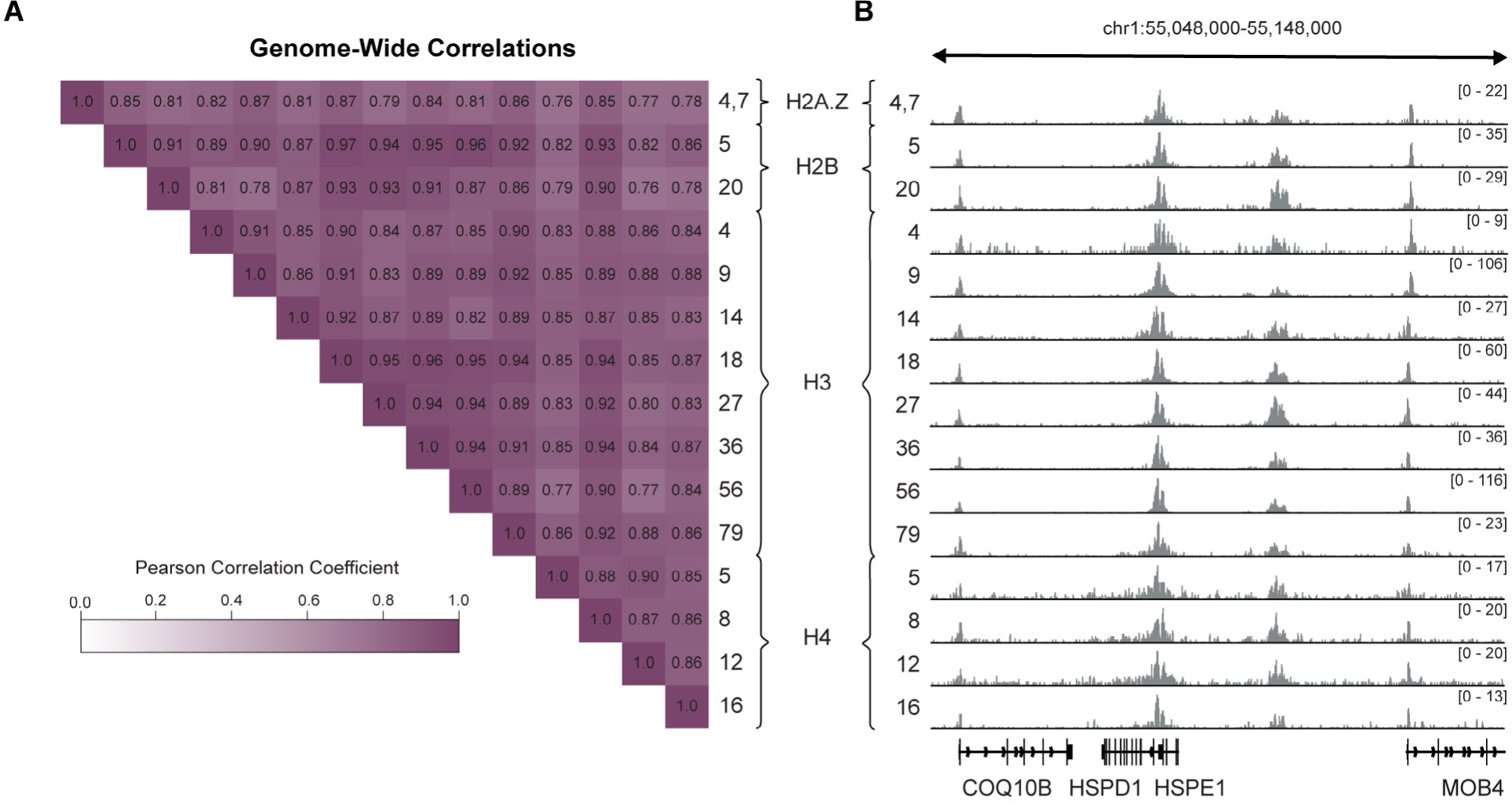
Histone acetylation marks are highly correlated genome-wide, related to Figure 7. **(A)** Genome-wide pearson correlation of 15 different histone acetylation marks in mESC. Correlations are based on coverage computed in 10kB windows. **(B)** Comparison of 15 different histone acetylation marks across a genomic region (mm10, chr1:55,048,000-55,148,000) in mESC.

**Supplemental Figure 7:**
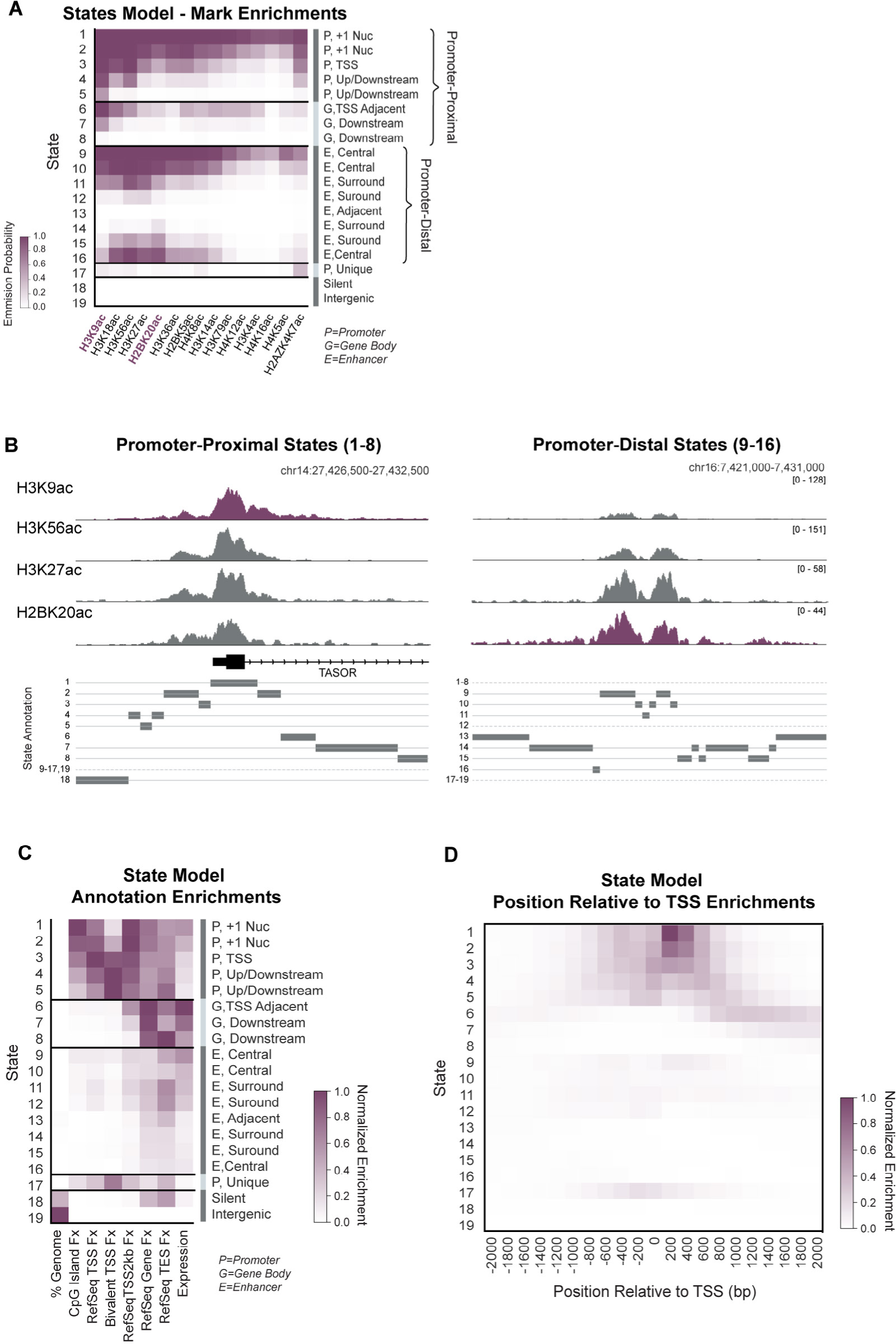
ChromHMM model using histone acetylation marks, related to Figure 7. **(A)** Histone acetylation mark emission probability matrix for 19-state ChromHMM model. State annotations (right) were assigned manually based on genomic position enrichments of states. **(B)** Track visualization of histone acetylation marks (top) and chromatin state annotations (bottom) at example promoter region (left) versus example intergenic region (right). Histone acetylation marks are scaled to the same maximum values at both regions. At each region, the chromatin states that are present are shown with solid lines and a box indicating the exact position; chromatin states that are absent are listed next to dotted lines. **(C)** Heatmap of genome annotation enrichment of chromatin states. Enrichment scores are normalized to the maximum and minimum of each column. **(D)** Heatmap of genomic position enrichment relative to the TSS of chromatin states. Enrichment scores are normalized to the maximum and the minimum of the heatmap.

**Supplemental Figure 8:**
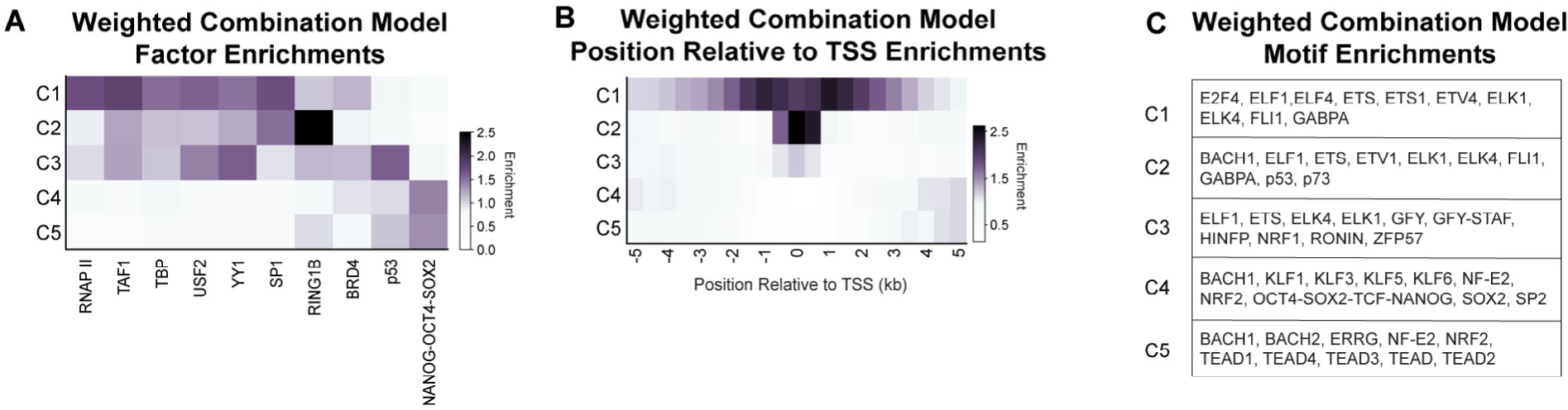
Enrichment profiles for NMF generated combinations (C1-C5) of histone acetylation marks, related to Figure 7. **(A)** RNAP II, TF and CR enrichment matrix for regions assigned to combinations (C1-C5) from NMF decomposition of highly acetylated regions using histone acetylation marks. **(B)** Heatmap of genome position enrichments relative to TSS for regions assigned to combinations. **(C)** Transcription factors of top 10 most significant sequence motifs for regions assigned to each combination are listed.

**Supplemental Figure 9:**
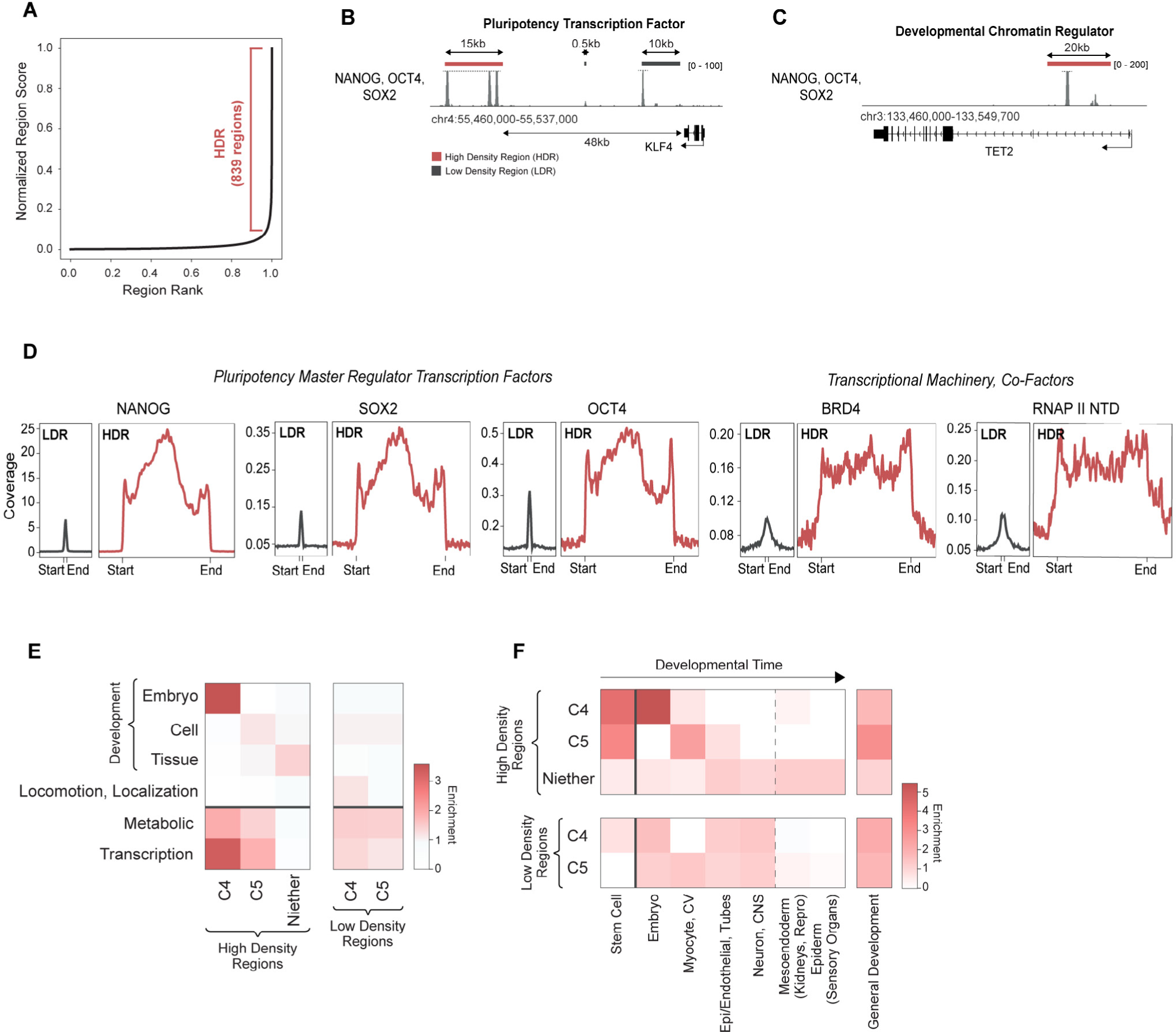
Profiles for high density regions of NANOG-OCT4-SOX2, related to Figure 7. **(A)** Plot showing normalized region scores (x-axis) for peak regions of NANOG-OCT4-SOX2, ordered by rank (y-axis). High density regions are defined as regions past the point where the slope = 1. **(B)** Track visualization of NANOG-OCT4-SOX2 upstream of the gene for KLF4, a pluripotency transcription factor, in mESC. A high density region is indicated with a red bar; low density regions are indicated with grey bars. (**C)** Visualization of NANOG-OCT4-SOX2 near the TET2 gene, a developmentally associated chromatin regulator, in mESC. A high density region internal to the gene is indicated with a red bar. **(D)** Coverage metaplots over low density regions (LDR) vs high density regions (HDR) for pluripotency transcription factors and other transcriptional-related factors. Metagenes are centered on the region and the lengths represent the approximate difference in mean lengths (500bps for LDRs and 14,500bps for HDRs). An additional 4kb surrounding each region is shown. **(E)** Enrichment heatmap for GO terms of genes associated with HDRs or LDRs containing C4, C5 or neither C4/C5 chromatin signatures. **(F)** Enrichment heatmap for development-associated GO terms of genes associated with HDRs or LDRs containing C4, C5 or neither C4/C5 chromatin signatures.

